# TPS Expression, Not Copy Number, Explains Aroma Diversity among Grapevine (*Vitis vinifera* L.) Cultivars

**DOI:** 10.64898/2026.08.03.742548

**Authors:** Malin Petersen, Manon Paineau, Andrea Minio, Noé Cochetel, Rosa Figueroa-Balderas, Lars H. Kruse, Sarah K. Davis, Jörg Bohlmann, Dario Cantu, Simone D. Castellarin

**Affiliations:** Wine Research Centre, Faculty of Land and Food Systems, The University of British Columbia, Vancouver, British Columbia V6T 1Z4, Canada; Department of Viticulture and Enology, University of California Davis, CA 95616, USA; Michael Smith Laboratories, University of British Columbia, Vancouver, British Columbia V6T 1Z4, Canada; Department of Biology, Concordia University, Montreal, QC, Canada; Department of Botany, University of British Columbia, Vancouver, British Columbia V6T 1Z4, Canada; Department of Forest and Conservation Science, University of British Columbia, Vancouver, British Columbia V6T 1Z4, Canada; UBC Botanical Garden, University of British Columbia, Vancouver, British Columbia, V6T 1Z4, Canada; Genome Center, University of California Davis, CA 95616, USA

## Abstract

Terpenoids are major contributors to grapevine berry and wine aroma, with mono- and sesquiterpenoids exhibiting substantial variation among cultivars. However, the genetic basis of cultivar-specific terpenoid profiles remains only partially understood due to gene duplications that expanded the terpenoid synthase (TPS) gene family across different cultivars. Here, we compared haplotype-resolved genome assemblies for seven cultivars to analyze copy number variation and expression profiles of terpenoid biosynthesis genes and terpenoid accumulation throughout flower and berry development. These assemblies resolved duplicated *TPS* loci at the haplotype level, revealing extensive copy number variation across cultivars and between haplotypes within a cultivar, with up to 44 mono-*TPS* genes on a single haplotype. We identified eight specific monoterpenoids as key drivers of cultivar-specific profiles in ripe berries. Total *TPS* transcript abundance correlated strongly with total terpenoid accumulation over time. In contrast, *TPS* copy number was largely decoupled from both expression and terpenoid accumulation, indicating that expression regulation, rather than gene dosage, underlies aroma differences among cultivars. Transcript-metabolite network analysis revealed both expected correlations between specific *TPS* genes and putative products as well as novel associations. We functionally validated selected candidate *TPS* in *Nicotiana benthamiana*, characterizing *α*-farnesene, *β*-ocimene, and multi-product *α*-terpineol synthases. The *in vivo* characterization demonstrated that minor sequence variations can alter catalytic activity and product profile. Together, these findings show that the extensive duplication of *TPS* genes in grapevine does not itself determine terpenoid output; instead, cultivar-specific aroma arises from how these genes are expressed and from sequence differences that shape enzyme activity.

## 3 Introduction

Terpenoids are a diverse group of specialized (i.e., secondary) metabolites that play important roles in many different chemo-ecological interactions of plants with other organisms. Mono- and sesquiterpenoids are made up of two or three isoprenoid units, respectively, and are major contributors to grapevine (*Vitis vinifera* L.) berry and wine aroma (Lund and Bohlmann, 2006). The complex combination of mono- and sesquiterpenoids at harvest determines cultivar-specific floral, fruity, and herbal aroma of berries, and is a major driver of economic value of derived wines (Carrascosa et al., 2001). The plastidial methyl-erythritol-phosphate (MEP) and the cytosolic mevalonic acid (MVA) pathways provide the substrates GPP and FPP for mono- and sesquiterpenoid biosynthesis, respectively (Lichtenthaler et al., 1997; Rodrıguez-Concepción and Boronat, 2002). 1-deoxy-D-xylulose 5-phosphate synthase (*DXS*), the first gene in the MEP pathway, is a key driver of metabolic flux of plastidial terpenoid biosynthesis (Tholl, 2015). In *V. vinifera*, quantitative trait loci (QTL) containing *DXS* genes have been linked to terpenoid accumulation (Battilana et al., 2009; Duchêne et al., 2009). The isoprenoid products of the MEP and MVA pathway are utilized by prenyl transferases to form the linear 10-, 15- and 20-prenyl diphosphate substrates for terpenoid synthases (TPS). The *TPS* genes occur in grapevine as a large gene family, including members of subfamilies TPS-a, TPS-b, TPS-c, TPS-e/f, and TPS-g. Subfamilies TPS-a, TPS-c, and TPS-e/f mostly contain sesqui- and di-TPS, whereas TPS-b and TPS-g contain mono-TPS. Due to their importance in berry aroma, over 40 *TPS* genes have been functionally characterized in grapevine (Drew et al., 2016; Ilc et al., 2017; Martin et al., 2010; Martin and Bohlmann, 2004; Matarese et al., 2013; Zhu et al., 2014).

Despite the relevance of grape berry aroma for determining wine quality and economic value, and the extended scientific effort invested in the study of these aromas, the genetic aroma potential of cultivars had not been explored in much detail until recently (S. Liu et al., 2022). The sequencing and various assemblies of the reference genome PN40024 (Canaguier et al., 2017; Jaillon et al., 2007; Shi et al., 2023; Velt et al., 2023) demonstrated that the *TPS* gene family is highly expanded in grapevine, with 69 putatively functional *TPS* compared to 32 in *Arabidopsis thaliana* (Aubourg et al., 2002; Drew et al., 2016; Lücker et al., 2004; Martin et al., 2010; Martin and Bohlmann, 2004; Smit et al., 2019; Zhu et al., 2014). Recent studies have shown significant differences in *TPS* gene copy numbers and expression among newly sequenced genotypes and proposed a key role in terpenoid accumulation (Bosman et al., 2023; Lin et al., 2026; Smit et al., 2020). However, there is still limited knowledge regarding how this genomic structural variation and possible differences in terpenoid biosynthesis gene expression between wine grape cultivars influence terpenoid accumulation patterns throughout development. Addressing this question has also been constrained by the available genomic resources. Haplotype-resolved reference genomes are required to understand the repeatedly duplicated and heterozygous *TPS* loci (Lin et al., 2026). However, these assemblies are largely limited to low-terpenoid (non-aromatic) cultivars, while high-terpenoid (aromatic) cultivars relevant to terpenoid diversity are largely lacking at this resolution.

In this study, we assembled four diploid genomes (Albariño, Fiano, Gewürztraminer, and Viognier) and used three previously available diploid genomes (Cabernet Sauvignon, Pinot Noir, and Riesling) to enable haplotype-resolved identification of copy number variation (CNV) in terpenoid biosynthesis genes, including MEP pathway, MVA pathway, prenyl transferases, and *TPS* genes via manual annotation. We hypothesized that *TPS* gene copy number differs between haplotypes and cultivars due to differential duplication and post-duplication events and examined whether this structural variation contributes to differences in grape aroma among cultivars. To investigate cultivar-specific regulation, the expression of identified terpenoid biosynthesis genes across haplotypes was analyzed using RNA-Seq at anthesis and throughout berry development (Figure 1).

**Figure 1:**
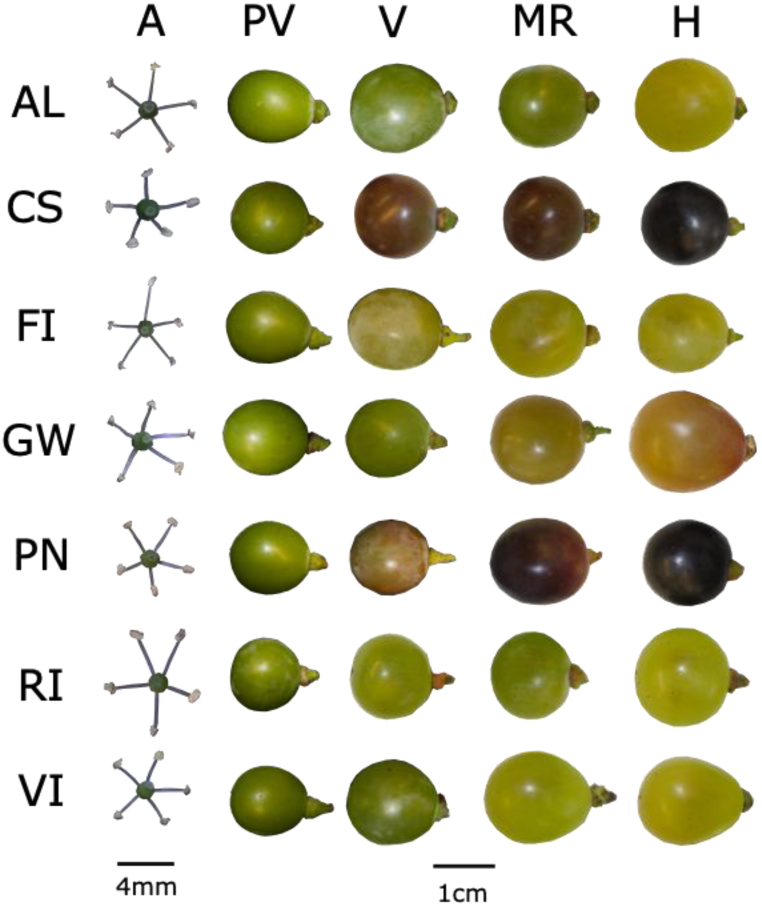
Flowers and berries of Albariño (AL), Cabernet Sauvignon (CS), Fiano (FI), Gewürztraminer (GW), Pinot Noir (PN), Riesling (RI), and Viognier (VI) at five sampling stages: Anthesis (A), pre-veraison (PV), veraison (V), mid-ripening (MR), and harvest (H).

These transcriptomic profiles were correlated with free (i.e., terpenoids that were not conjugated to sugars) and bound (i.e., terpenoids conjugated to sugars) terpenoid concentrations from flowers and developing berries. Selected *TPS* genes were functionally characterized representing orthogroups that lacked validated functional information using heterologous expression in *Nicotiana benthamiana*.

## 4 Materials and Methods

### a. DNA extraction and sequencing libraries for genome assemblies

The previously published Cabernet Sauvignon clone FPS 08 (CS), Pinot Noir clone FPS 123 (PN), and Riesling clone FPS 24 (RI) genomes were used for genomic and transcriptomic analyses (Cochetel et al., 2025a; Lin et al., 2026; Paineau et al., 2025). Leaf material of Albariño clone FPS 03.1 (AL), Fiano clone FPS 2.1 (FI), Gewürztraminer clone FPS 01 (GW), and Viognier clone FPS 05 (VI) vines was available through the Foundation Plant Services (FPS) at the University of California, Davis, and high-quality genomic DNA was isolated following the protocol described in Chin et al. (2016). HiFi libraries were prepared for Gewürztraminer using SMRTbell® Express Template Prep Kit 2.0 and subsequent treatment with the Enzyme Clean Up Kit (Pacific Biosciences,Menlo Park, CA, USA) The HiFi SMRTbell® templates were size selected using the BluePippin system (Sage Sciences, Beverly, MA, USA), and the final libraries were purified using AMPure PB beads (Pacific Biosciences, Menlo Park, CA, USA). Qubit™ 1X dsDNA HS Assay Kit (Thermo Fisher, Waltham, MA, USA) and Femto Pulse System (Agilent, Santa Clara, CA, USA) were used to determine concentration and library size distribution. Continuous long read (CLR) libraries were prepared for Albariño, Fiano, and Viognier following Cochetel et al. (2023). HiFi and CLR libraries were sequenced on a PacBio Sequel II platform (DNA Technology Core Facility, University of California, Davis, CA, USA).

### b. Genome assembly, gene prediction, functional annotation and terpenoid biosynthesis gene identification

All four newly generated genomes were produced through a two-step assembly strategy. The initial scaffolding step relied on the sequencing technology. HiFi reads from Gewürztraminer were assembled with Hifiasm v0.16.1-r37412 (Cheng et al., 2022, 2021), whereas CLR reads from Albariño, Fiano, and Viognier were processed with the FALCON-Unzip pipeline (v.2017.06.28-18.01; Chin et al., 2016). To obtain minimally fragmented draft assemblies, the Gewürztraminer HiFi dataset was assembled with Hifiasm following the procedure described in Paineau et al., (2025) using the parameters a=4, k=35, w=41, f=0, r=3, s=0.6, D=4, N=25, n=9, and m=2,500,000. For the Albariño, Fiano, and Viognier, scaffolding of CLR reads was carried out according to the workflow reported by Minio et al. (2019a). The customized FALCON-Unzip pipeline is available at https://github.com/andreaminio/FalconUnzip-DClab.

In a second step, haplotype-resolved chromosome-scale pseudomolecules were reconstructed for each genome using the HaploSync tool suite v1.0 (Minio et al., 2022) in combination with the grapevine high-density gene map (Cochetel et al., 2025b) for Albariño, Fiano, and Viognier and the consensus grape genetic map (Zou et al., 2020) for Gewürztraminer. Assembly metrics are provided in Table S1.

Gene structural annotations were generated using the workflow described by Minio et al. (2019b). Briefly, gene prediction relied on a combination of ab initio tools, including BUSCO v3.0.2 (Waterhouse et al., 2018), Augustus v3.0.3 (Stanke et al., 2006), GeneMark v3.47 (Lomsadze et al., 2005), and SNAP v2006-07-28 (Korf, 2004). All predictors were parameterized using a training model previously developed for grapevine (Massonnet et al., 2020). Repetitive sequences were first identified and masked with RepeatMasker v4.0.6 (Smit et al., 2013) using a curated *Vitis vinifera* repeat library (Minio et al., 2019b). Evidence from transcript and protein datasets described in Minio et al. (2019b) was then aligned to the genomes using PASA v2.3.3 (Haas et al., 2003), MagicBLAST v1.4.0 (Boratyn et al., 2019), and Exonerate v2.2.0 (Slater and Birney, 2005). The outputs from ab initio predictions and alignment-based evidence were consolidated with EvidenceModeler (EVM) v1.1.1 (Haas et al., 2008) to produce consensus gene models. Predicted genes containing internal stop codons were discarded. Functional annotation of the final gene set followed the approach reported by Cochetel et al. (2021). Genome completeness was evaluated using BUSCO v5 (Benchmarking Universal Single-Copy Orthologs; Simão et al., 2015) with the ‘viridiplantae_odb10’ datasets (425 orthologs). Analyses were performed separately and together for haplotypes 1 and 2 as well as for the unplaced contigs (Table S1). Telomeric motifs were investigated with TIDK v0.2.63 (Brown et al., 2025). The ‘tidk explore’ function was applied to the terminal 1% of each chromosome to identify candidate repeat units between 5 and 12 bp, requiring at least two consecutive occurrences. The telomeric repeat sequence TTTAGGG was then searched using ‘tidk search’.

Genes previously described as involved in terpenoid biosynthesis, including MEP and MVA pathway genes and prenyl transferases were selected from TAIR (for *A. thaliana;* https://www.arabidopsis.org/) and KEGG pathway databases (*Solanum lycopersicum* L. https://www.kegg.jp/pathway/sly00900; *A. thaliana* https://www.kegg.jp/pathway/ath00900; and *V. vinifera* https://www.kegg.jp/pathway/vvi00900). Functionally characterized *V. vinifera TPS* genes were identified from previous publications (Martin et al., 2010; Martin and Bohlmann, 2004; Matarese et al., 2013; Zhu et al., 2014; Drew et al., 2016; Lücker et al., 2004). These MEP and MVA pathway, prenyl transferases, and *TPS* genes were used to identify terpenoid biosynthesis genes in the original PN40024v1 and PN40024v4.3 reference genome, which has higher contiguity, as well as in Albariño, Cabernet Sauvignon, Fiano, Gewürztraminer, Pinot Noir, Riesling, and Viognier genomes via a four-step process: (1) BLASTN: alignment of identified genes as queries to each genome (e-value <1e^-10^) (Camacho et al., 2009); (2) GMAP: alignment of identified transcripts and BLASTN matches to all genomes (percentage identity >80%) (Wu et al., 2016; Wu and Watanabe, 2005); (3) PFAM: curation of the terpenoid biosynthesis genes set built by BLASTN and GMAP with searches for hidden Markov model (HMM) matches (Eddy, 1996). For *TPS* gene identification, PF01397 and PF03936 motifs in the N- and C-terminal of the protein were used, respectively (Martin et al., 2010; Smit et al., 2020, 2019; Zhu et al., 2014). To maximize the capture of potential *TPS* genes, all genes with an alignment length to one PFAM motif >50 amino acids (Martin et al., 2010), a total amino acid sequence of 350 or longer (Bohlmann et al., 1998; Durairaj et al., 2019), and more than 4 exons (Smit et al., 2020) were considered putatively functional *TPS* genes. For the HMM analysis of MEP and MVA pathway genes and prenyl transferases the following PFAM domains were used: *AACT*: PF02803, PF00108; *HMGS*: PF01154, PF08540; *HMGR*: PF00368; *MVK*: PF00288, PF08544; *PMVK*: PF00288, PF08544; *MVD*: PF00288, PF18376; *FPPS*: PF00348; *DXS*: PF02779, PF02780, PF13292; *DXR*: PF02670, PF08436, PF13288; *MCT*: PF01128; *CMK*: PF00288, PF08544; *MECPS*: PF02542; *HDS*: PF04551; *HDR*: PF02401; *GPPS*: PF00348; *IPPI*: PF00293. Pathway genes were considered putatively functional when their PFAM motif alignment length fell within ±2 standard deviations (SD) of the characterized genes or, if the SD was 0, within 10% of the reference alignment length (Durairaj et al., 2019). Previous *DXS* and *HMGR* classifications were assigned based on sequence similarities (Leng et al., 2017). (4) Phylogenetic analysis: protein sequences were globally aligned using the MAFFT (v7.511) multiple sequence alignment tool (Katoh et al., 2002) and maximum likelihood phylogenetic trees generated using the RAxML-NG tool (v. 0.9.0), 1000 bootstrap replicates with BLOSUM62 mode activated (Kozlov et al., 2019). FigTree (version 1.4.4) was used for tree visualization (http://tree.bio.ed.ac.uk/software/figtree/). *TPS* subfamilies and subclades of pathway genes were assigned based on phylogenetic clades with characterized genes. Orthofinder v. 2.3.7 was used to identify orthologs between *TPS* genes of different cultivars and paralogs within haplotypes (Emms and Kelly, 2019, 2015). Presence of characterized genes in the orthogroups (OG) was used to assign putative function to members. The OGs were divided into 3 categories: core, dispensable, and private. Core groups contained at least one gene for each genome, dispensable groups contained genes on multiple, but not all genomes, and private genes were only found in one genome. Subcellular localization of TPS proteins was predicted using TargetP based on the presence of N-terminal transit peptides (Armenteros et al., 2019).

### c. Plant material for transcriptomic and metabolomic analyses

Plant material for transcriptomic and metabolomic analysis was collected in 2023 from the same clones mentioned above except for Cabernet Sauvignon, which used clone FPS 04. Samples were represented by four biological replicates (three for Fiano at pre-veraison and harvest due to material limitation) and were collected at five developmental stages: anthesis (A, EL23, 50 % anthesis), pre-veraison (PV, EL31, pea-sized green berry), veraison (V, EL35, 50 % veraison), mid-ripening (MR, EL36, 18-19 °Brix), and harvest (H, EL38, 21-22 °Brix) (Figure 1).

Due to differences in phenology between cultivars, total soluble solids (TSS) were used as a proxy for the ripening stage of berries. Thirty berries were collected for immediate basic berry parameter analysis including TSS, pH, and titratable acidity (TA). The clarified juice of these 30 berries was used to measure TSS expressed as °Brix via a Atago, PR-101α palette digital refractometer, pH via a Mettler Toledo LE409 pH meter, and TA via acid-base titration as previously described by Kovalenko et al. (2021). The TSS levels at each developmental stage were kept within 1 °Brix difference between means of cultivars for all but the veraison stage, at which the difference between means was up to 2.5 °Brix. Differences in TSS among cultivars at each developmental stage were assessed using one-way ANOVA (Figure S1).

Sampling for all stages occurred between 6 and 9 am. Twenty flowering clusters or 40-45 berries were randomly collected for each replicate at anthesis and berry sampling, respectively. Flowering clusters were removed with scissors at the stem and berries at the pedicel to avoid damage to the berry. Flowers were separated from rachises and flash frozen in liquid nitrogen. Berries were placed on ice, seeds and rachises were removed and they were flash frozen immediately. The frozen flowers and berries were ground to a fine powder using a Retsch Mixer Mill 400 for 30 seconds with a frequency of 30 Hz. The powder was stored at −80 °C until further processing.

### d. RNA extraction and sequencing

Total RNA extraction and library preparation were performed on flowers and berries following Rapicavoli et al. (2018) and Blanco-Ulate et al. (2013), respectively.

RNA purity was assessed with a NanoDrop 2000 spectrophotometer (Thermo Scientific, Hanover Park, IL, USA), RNA concentration with a Qubit 2.0 Fluorometer and Broad Range RNA kit (Life Technologies, Carlsbad, CA, USA), and RNA integrity through agarose gel electrophoresis. RNA-Seq libraries were evaluated using an Agilent 2100 Bioanalyzer (Agilent Technologies, CA, USA). Sequencing was conducted on an Aviti sequencer at the DNA Technology Core Facility, University of California, Davis, CA, USA, generating 75-bp paired-end reads. On average 48.32 ± 0.92 million reads (mean ± SE) were obtained per sample (Table S2).

The reads were checked for quality and the adaptor sequences removed using fastQC (v0.11.5; Babraham Bioinformatics) and trimmomatic (v0.36; Bolger et al., 2014), respectively. Clean reads were mapped and quantified using Salmon (v1.4.0). Sequence-specific and GC-content biases were corrected using the ‘--seqBias’ and ‘--gcBias’ options, respectively (Patro et al., 2017). On average 89.61 ± 0.36% of the reads were mapped to the reference genomes and the mean GC content was 46.02 ± 0.05% (Table S2). Read counts were extracted using the tximport package in R (Soneson et al., 2016). DESeq2 (v1.46.0) was used to normalize reads using the ‘normalize=TRUE’ parameter (Anders and Huber, 2010; Zhao et al., 2021) and create a distance matrix and PCA plot to identify potential outliers; however none were detected (Love et al., 2023, 2014). Log_10_-transformed expression of terpenoid biosynthesis genes was used for visualization and correlation analysis (Zhao et al., 2021). The expression of MEP and MVA pathway genes, prenyl transferases, and OGs, summarized in Table S5-6, was calculated by totaling the expression of genes with the same putative functions.

### e. Profiling of free and bound terpenoids

Free (i.e., terpenoids that were not conjugated to sugars) and bound (i.e., terpenoids conjugated to sugars) terpenoid analysis was carried out using SPME-GC/MS as previously described by Pico et al. (2022, 2023), respectively. The bound terpenoids were released from sugars by pectinase AR2000^®^ (Sigma Aldrich) for analysis, as described in Pico et al. (2023). As bound terpenoids, we included glycoside esters as well as alcohols and acids released from conjugates. Other compounds were deemed potential artifacts of the extraction method (Pico et al. 2023), and not included in the analysis. A 7890A gas chromatograph and 5975C single quadruple mass spectrometer (GC/MS) were coupled to a GC 80 autosampler with headspace-solid phase microextraction (SPME) and Chemstation software (version E.02.02.1431, Agilent Technologies, Santa Clara, CA, USA). Quantification of free and bound terpenoids was achieved with an internal standard, external calibration curves, and matrix effect. For the calibration curves 100 μL of standard mix of terpenoids was used to prepare samples as described in Pico et al. (2022, 2023). The matrix effect was calculated independently for each developmental stage and for red and white cultivars separately using the following equation:

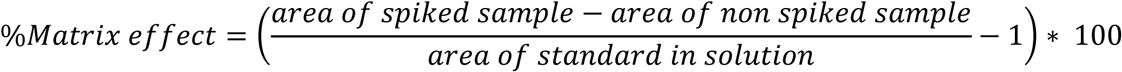

Limit of detection (LOD) and limit of quantification (LOQ) were calculated following Pico et al. (2023). The final concentrations (µg kg^-1^ fresh weight) were used for visualization and correlation analysis.

### f. Transient expression and functional characterization of *TPS* candidates in *N. benthamiana*

*TPS* coding sequences, including their native transit peptides for subcellular targeting, were synthesized (Twist Bioscience, San Francisco, CA, USA). Internal BpiI, BsaI, and Esp3I recognition sites were eliminated by synonymous substitutions to enable Golden Gate-based Modular Cloning (MoClo; Engler et al., 2014; Weber et al., 2011). Flanking adaptor sequences containing BpiI recognition sites were incorporated to generate phytobrick-compatible Level 0 modules. The coding sequences were assembled into the Level 0 acceptor vector pICH41308 (Addgene #47998) by BpiI-mediated Golden Gate assembly, resulting in phytobrick CDS modules. Constructs were transformed into NEB 5-alpha competent *Escherichia coli*, and positive clones were confirmed by colony PCR and sequencing. For level 1 assembly, *TPS* and *GPPS* CDS modules were excised using flanking BsaI sites and assembled into the destination vector pICH47732 (Addgene #48000) via BsaI-mediated Golden Gate cloning. The level 1 backbone contains the cauliflower mosaic virus 35S promoter (P35S), the 5′ untranslated region of barley stripe mosaic virus (pICH44188), and the *Agrobacterium tumefaciens* mas 3′ untranslated region and polyadenylation signal (pICH77901). *DXS* and *HMGR* were cloned with the USER cloning as described in Kruse et al. (2024). In brief, complete open reading frames for each gene were cloned into the pCAMBIA230035Su vector following the USER cloning method (Geu-Flores et al., 2007; Nour-Eldin et al., 2006; Table S3).

*N. benthamiana* plants were grown in a Conviron (ARC10) growth chamber under 16h light/8h dark photoperiod at 22 °C, and a light intensity of 125 µmol m⁻² s⁻¹. Competent *A. tumefaciens* cells strain GV3101pMP90 were transformed with binary plasmids (pCAMBIA230035Su or pICH47732) containing the gene of interest (Koncz and Schell, 1986). Single colonies were inoculated into lysogeny broth medium supplemented with antibiotics as appropriate: 25 µg mL⁻¹ rifampicin and 25 µg mL⁻¹ gentamicin (GV3101 background) plus either 50 µg mL⁻¹ carbenicillin (pICH47732) or 50 µg mL⁻¹ kanamycin (pCAMBIA230035Su). Cultures were grown overnight at 28 °C at 210rpm. Cells were harvested by centrifugation at 4,300 rpm for 10 min at room temperature and resuspended in infiltration buffer (10 mM 2-(N-morpholino)ethanesulfonic acid [MES], pH 5.7; 10 mM MgCl₂; 100 µM acetosyringone). Suspensions were adjusted to an OD₆₀₀ of 0.5 and incubated for 2 h at room temperature with gentle agitation. In three co-expression experiments, (1) *TPS*, *GPPS*; (2) *TPS*, *DXS*, *GPPS*; (3) *TPS*, *DXS*, *GPPS*, *HMGR*, the *pBIN:p19 plasmid*, carrying the *p19* silencer of repression, was included in all infiltrations and cultures were combined in equal volumes. Two fully expanded leaves of 4-week-old plants were infiltrated in triplicate on the abaxial surface using a needleless syringe. Plants were maintained under the growth conditions described above for 5 days post-infiltration. Six leaf discs (12 mm diameter) were excised from infiltrated areas and immediately placed into 20 mL SPME vials. Vials were returned to the growth chamber for 24 h prior to headspace sampling. Products of transient expression trials were analyzed as in Pico et al. (2022), and identification of free terpenoids was achieved by comparison with retention times and dominant ions (Target, Q1, Q2, Q+) of pure standards. Chromatographic peak areas were compared to control samples infiltrated with only pathway and *p19* silencing suppressor genes.

### g. Statistical analysis

Data visualization and statistics were carried out using the R software (version 2023.06.1+524; R Core Team, 2024) Free and bound terpenoid concentrations and terpenoid biosynthesis gene expression were analyzed by one-way analysis of variance (ANOVA) and Tukey HSD test with p-adjusted<0.05 as the significance threshold. Only samples with values >0 for at least 2 replicates were considered. The analyses were carried out to identify significant differences between cultivars within a developmental stage and between different stages within one cultivar. The data was expressed as mean ± standard error (SE) and group letters were assigned with the multcompView R package (Donoghue, 1998; Piepho, 2004). Pearson and Spearman correlations were run with the stats package in R and transcript-metabolite and transcript-transcript correlation networks were visualized using the Cytoscape software (v3.10.1).

## 5 Results

### a. Haplotype-resolved genome assemblies and annotations

To investigate how terpenoid biosynthesis genes contribute to grapevine aroma, we analyzed chromosome-scale, haplotype-resolved genome assemblies of seven *V. vinifera* cultivars. Previously published genomes were used for Cabernet Sauvignon (Cochetel et al., 2025b), Pinot Noir (Paineau et al., 2025), and Riesling (Lin et al., 2026). In addition, we generated four new *de novo* assemblies for Albariño, Fiano, Gewürztraminer, and Viognier cultivars for which no haplotype-resolved reference was previously available.

Draft assemblies reflected the expected differences between HiFi- and CLR-based sequencing technology (Table S1). The HiFi assembly of Gewürztraminer directly produced two highly contiguous haplotypes (506.4 Mb and 501.4 Mb; contig N50 ∼15.1 Mb and ∼14.3 Mb, respectively), whereas CLR assemblies of Albariño, Fiano, and Viognier generated large primary contigs (mean 727.8 ± 4.4 Mb) accompanied by haplotigs (mean 183.2 ± 16.9 Mb) with substantially lower contiguity (contig N50 ∼2.3–2.4 Mb for primaries). After haplotype reconstruction and scaffolding with HaploSync, all 19 chromosomes were constructed per haplotype. The final pseudomolecule assemblies were highly consistent across cultivars, with a mean size of 412.5 ± 22.5 Mb per haplotype. Unplaced sequences ranged from ∼29.5 Mb in Gewürztraminer to ∼81.6 Mb in Viognier and were markedly enriched in repetitive elements (76–79%) compared with the anchored haplotypes (52–58%).

Telomeric repeat analysis indicated that assembly completeness was highest in Gewürztraminer, where 14 chromosomes per haplotype carried telomeric motifs at both ends. In the CLR-based assemblies, between 8 and 10 chromosomes per haplotype displayed telomeric repeats at both ends, with an additional 7 to 9 chromosomes carrying a single terminal repeat, supporting near-complete chromosome reconstruction in most cultivars.

Assembly quality was evaluated using the Benchmarking Universal Single-Copy Orthologs (BUSCO) framework (Simão et al., 2015), which indicated high completeness for all four haplotype-resolved assemblies, with scores exceeding 96% for each haplotype (range: 96.3– 99.0%). The entire genome assemblies, including unplaced scaffolds, contained 98.8–99.3% complete BUSCOs, with 94.6–96.9% classified as duplicated, consistent with the presence of both haplotypes. Unplaced scaffolds contained fewer than 5% of the detected BUSCO genes across all cultivars (range: 2.6–4.9%), supporting the overall integrity of the haplotype assemblies. Finally, gene annotation yielded between 27,958 and 29,475 protein-coding genes per haplotype depending on the cultivar (28,697 ± 561 genes).

### b. Largely expanded *TPS* gene families across haplotypes and cultivars

To investigate how CNV of terpenoid biosynthesis genes and their expression patterns impact terpenoid accumulation, we manually annotated these genes in the seven genomes of interest. Terpenoid biosynthesis genes identified as putatively functional are listed in Table S4. In this study CNV is based on comparative analysis of the number of terpenoid biosynthesis gene annotations between haplotypes and cultivars. CNV of MEP and MVA pathway genes and prenyl transferases was minor across haplotypes and cultivars, with the majority of genes being present in one copy per haplotype (Table S4).

*TPS* genes showed extensive CNV across cultivars and between the two haplotypes within a cultivar (Figure 2; Table S4).

**Figure 2:**
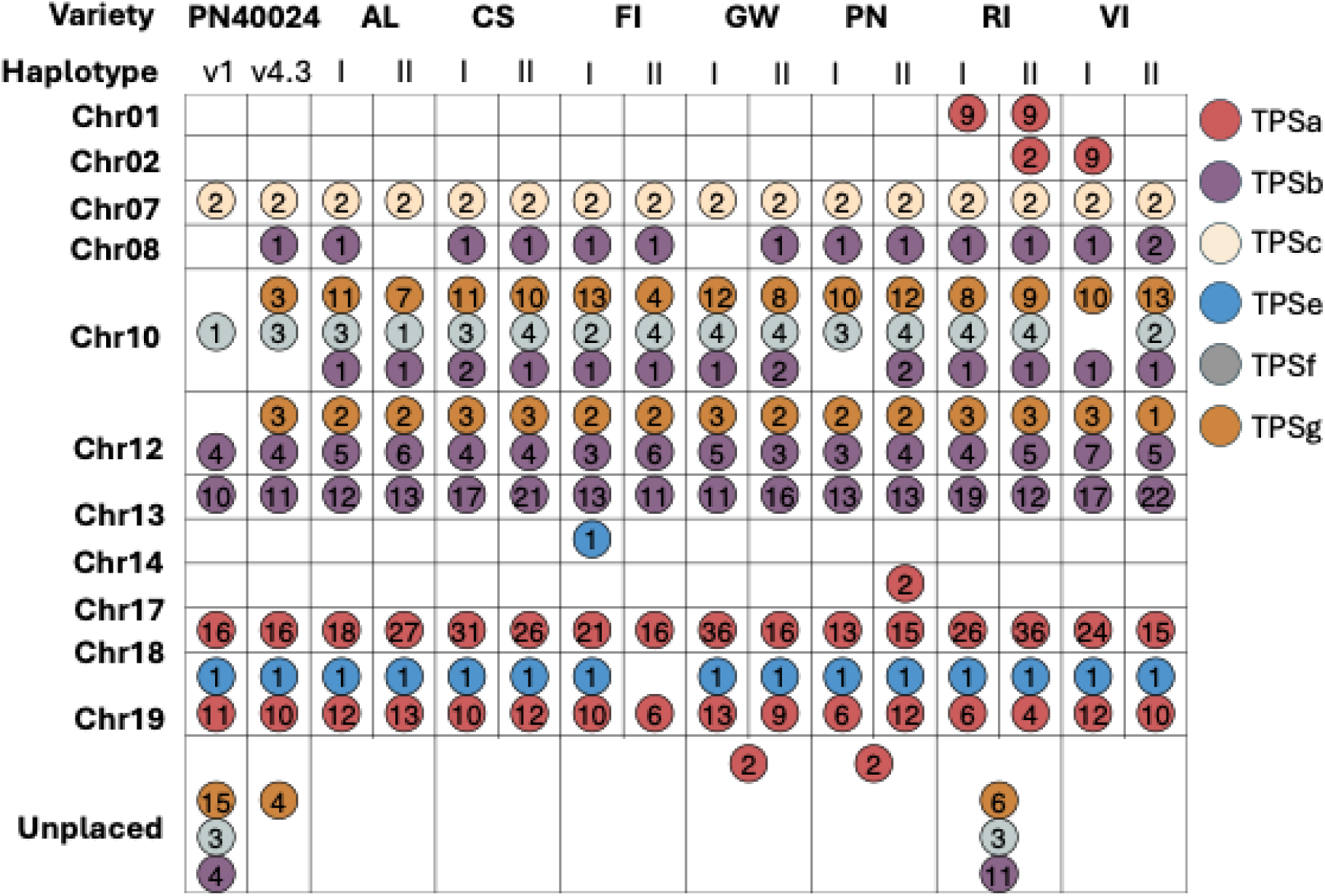
Diagram summarizing the TPS subfamily distribution across chromosomes for reference genomes PN40024v1 and PN40024v4.3 and haplotypes for each of the diploid genomes Albariño (AL), Cabernet Sauvignon (CS), Fiano (FI), Gewürztraminer (GW), Pinot Noir (PN), Riesling (RI), and Viognier (VI). The *TPS* subfamilies are colour-coded, and the numbers within circles represent the copy number of *TPS* genes within the given haplotype and chromosome, respectively. *TPS* genes that were on unplaced contigs are referred to as unplaced.

For instance, we identified 88 putatively functional *TPS* genes on haplotype I of Gewürztraminer compared to 64 on haplotype II (with 2 unplaced genes), whereas Pinot Noir exhibited 54 *TPS* on haplotype I and 68 on haplotype II (with 2 unplaced genes). Putative mono-*TPS* counts (subfamilies TPS-b and TPS-g) on a single haplotype ranged from 25 (Fiano) to 44 (Viognier), and putative sesqui-*TPS* (subfamily TPS-a) from 19 (Pinot Noir) to 51 (Riesling), and di-*TPS* (subfamily TPS-c, TPS-e, and TPS-f) from 3 (Viognier) to 7 (Cabernet Sauvignon, Gewürztraminer, Pinot Noir, and Riesling). The distribution of TPS subfamilies across chromosomes was conserved for all haplotypes and cultivars, and when multiple subfamilies were represented on one chromosome, their proportions were similar across haplotypes and cultivars (Figure 2). The mono-*TPS* in subfamilies TPS-b and TPS-g were located exclusively on chromosomes 8, 10, 12, and 13, with both chromosome 8 and 13 exclusively containing *TPS* genes from subfamily TPS-b. This subfamily-to-chromosome partitioning was conserved across all seven cultivars and both haplotypes of cultivars, indicating a stable ancestral organization of the mono-*TPS* array predating cultivar divergence. PN40024v1 and PN40024v4.3 contained notably fewer *TPS* genes on chromosome 10, and more without chromosome designation, represented by the unplaced genes, compared to other genomes (Figure 2). These results indicate that long read sequencing facilitated an improved assembly of *TPS* genes on chromosome 10 in the present genome assemblies compared to the PN40024 reference genomes.

Within *TPS* subfamilies we were able to assign orthologs and paralogs between cultivars and within haplotypes, respectively, using OrthoFinder (v.2.3.7) and subsequent manual curation based on phylogeny (Emms and Kelly, 2019, 2015; Table S4). This manual, phylogeny-based curation resolved 84 *TPS* OGs across the seven cultivars, of which 32 (12 mono-*TPS* and 20 sesqui-*TPS*) could be assigned putative functions through previously characterized members (Table 1; Figure 3-4; Figure S2-5).

**Figure 3:**
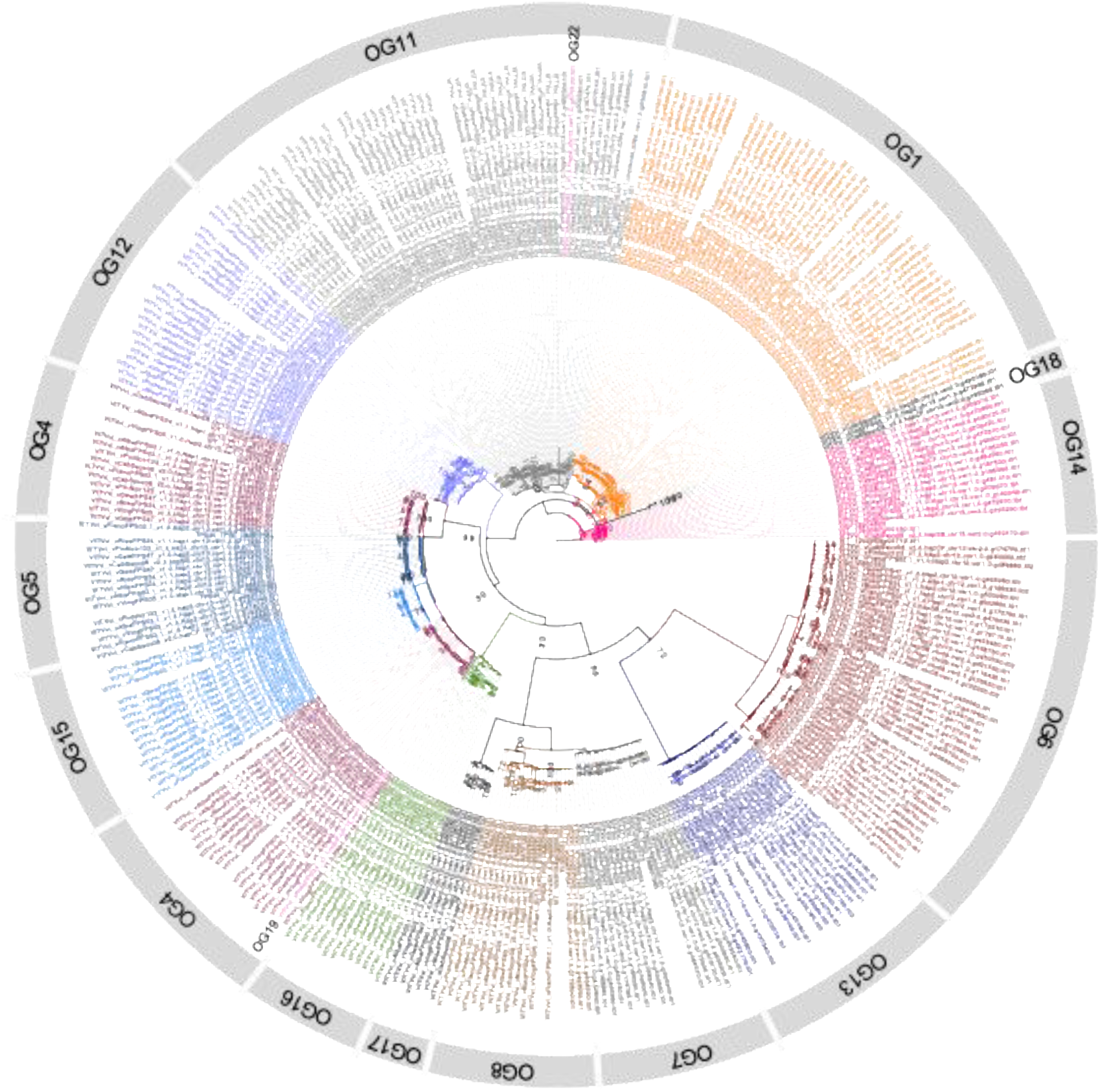
Phylogenetic tree (protein sequence) of *TPS* genes of subfamily TPS-b from Albariño, Cabernet Sauvignon, Fiano, Gewürztraminer, Pinot Noir, Riesling, Viognier, PN40024v1, PN40024v4.3, and characterized genes. Nodes are annotated with 1000 bootstrap values and clades are colour coded and annotated based on orthogroup (OG).

**Figure 4:**
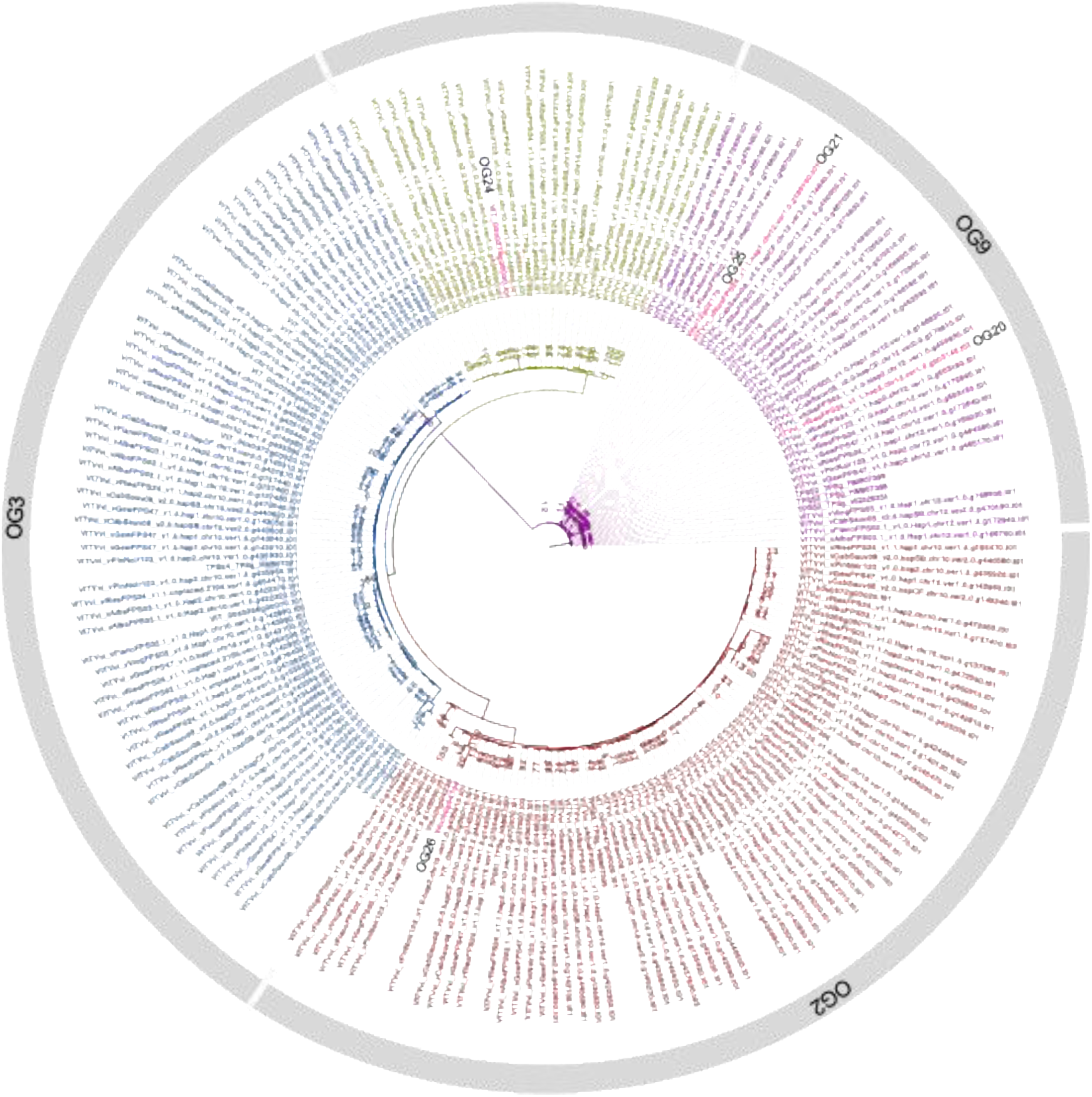
Phylogenetic tree (protein sequence) of *TPS* genes of subfamily TPS-g from Albariño, Cabernet Sauvignon, Fiano, Gewürztraminer, Pinot Noir, Riesling, Viognier, PN40024v1, PN40024v4.3, and characterized genes. Nodes are annotated with 1000 bootstrap values and clades are colour coded and annotated based on orthogroup (OG).

**Table 1:**
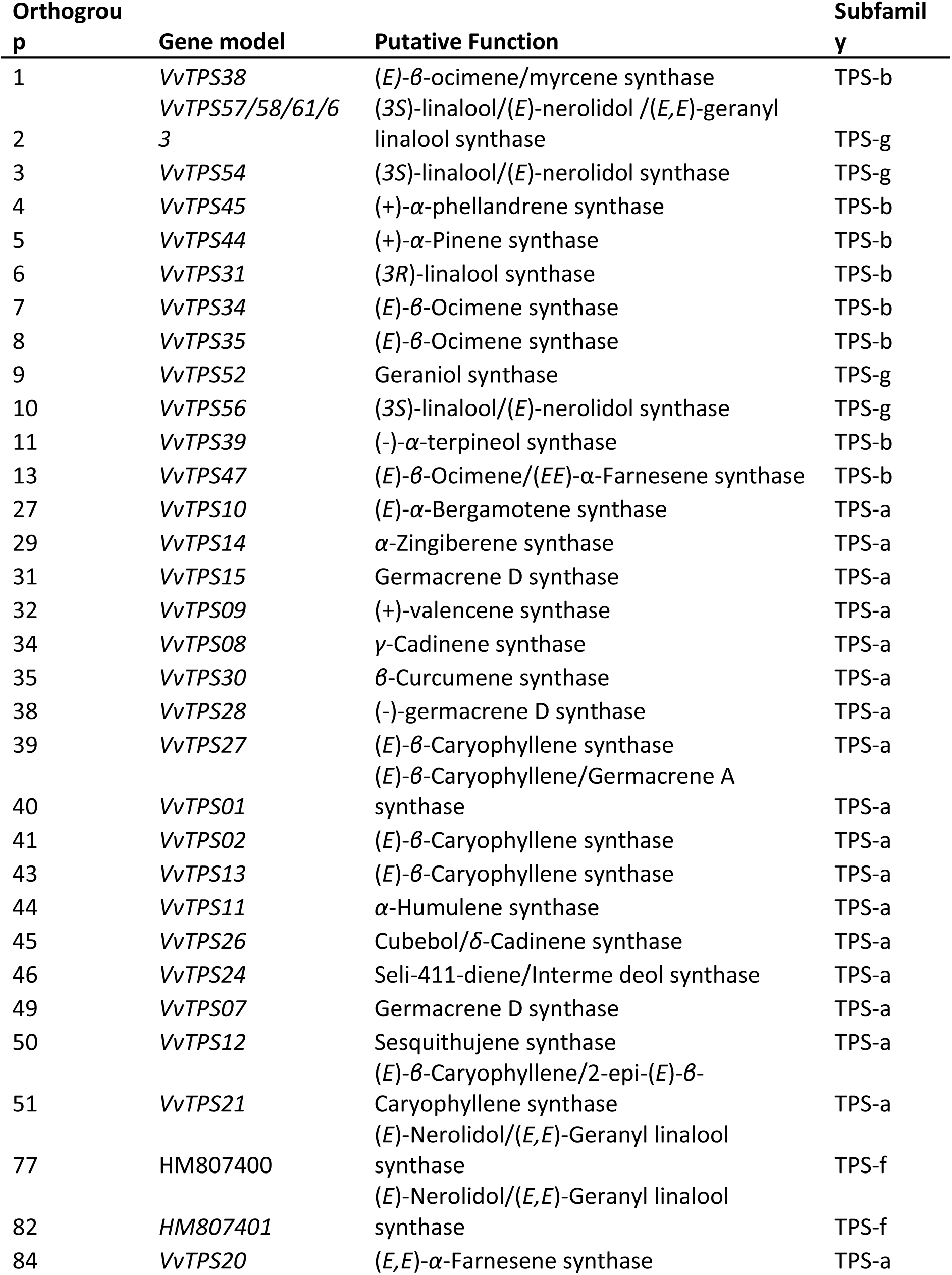
***TPS* OGs with putative functions based on closely related.**

Across cultivars *TPS* genes were present in clusters on chromosomes, comprised of multiple OGs (Table S4). However, gene CNV among cultivars differed substantially between OGs, ranging from two or fewer copies (OG5, OG6, OG7, OG9, OG12, OG16, OG17, and OG18) to 5 copies or more (OG1, OG4, OG11, OG14, and OG15).

### c. Transcriptomic analysis revealed different expression patterns of *TPS* genes across cultivars

To gain a deeper understanding of terpenoid biosynthesis, transcriptomes were analyzed via RNA-Seq. The mean expression levels of individual terpenoid biosynthesis genes throughout development were summarized in Table S4. Similar expression patterns were observed across all cultivars for both MEP and MVA pathway genes and prenyl transferases (Figure 5; Table S5-6).

**Figure 5:**
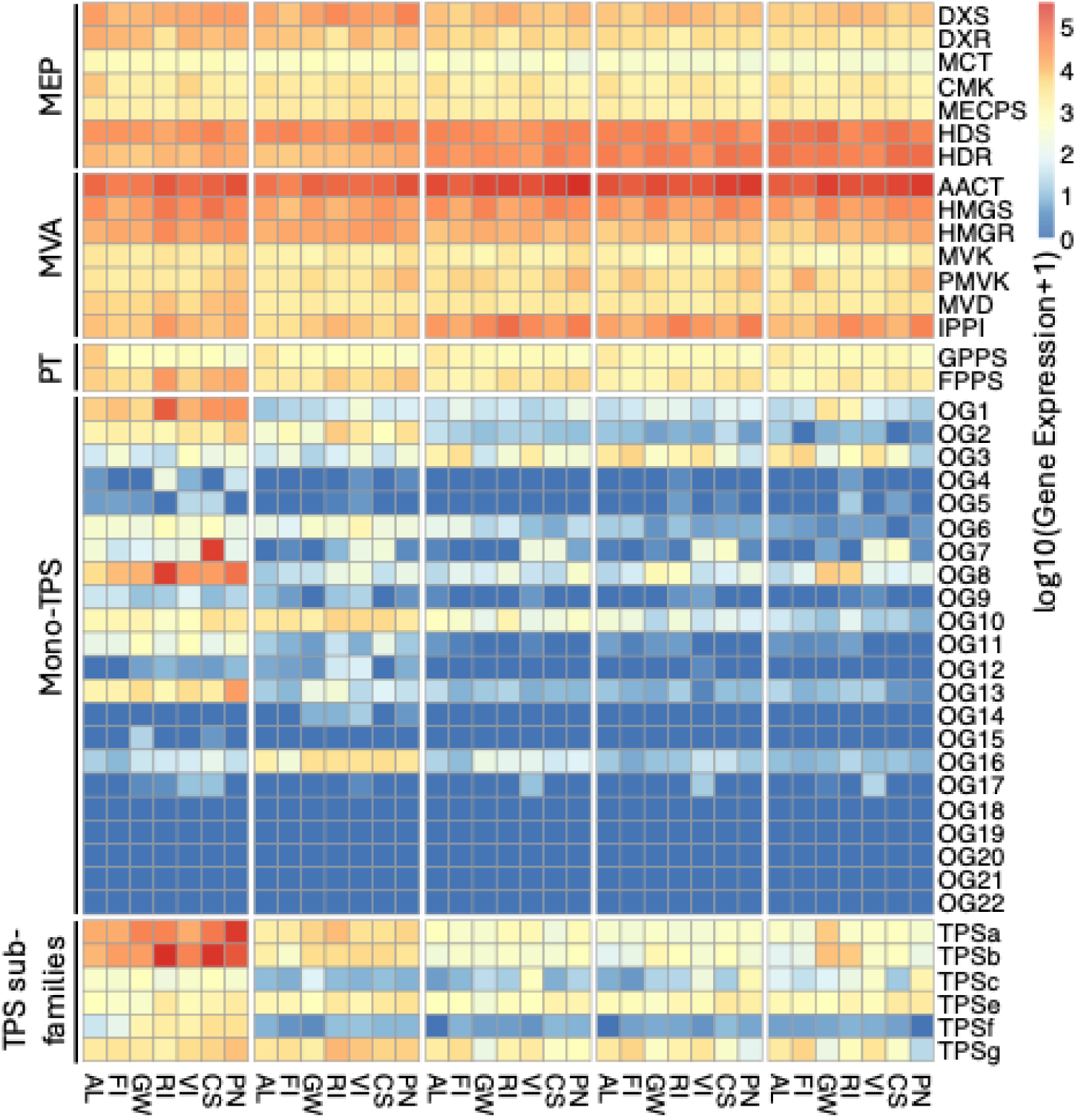
Heatmaps of log_10_-transformed total expression of MEP and MVA pathway genes, prenyl transferases, mono-TPS orthogroup genes, and total TPS subfamily expression throughout development. Strength of expression is colour coded and scaled across all terpenoid biosynthesis genes. Developmental stages are anthesis (A), pre-veraison (PV), veraison (V), mid-ripening (MR), and harvest (H) and the varieties are Albariño (AL), Cabernet Sauvignon (CS), Fiano (FI), Gewürztraminer (GW), Pinot Noir (PN), Riesling (RI), and Viognier (VI). Gene names are abbreviated functions of MEP pathway genes: *DXS*: 1-deoxy-dxylulose-5-phosphate synthase; *DXR*: 1-deoxy-D-xylulose-5-phosphate reductoisomerase; *MCT*: 4-diphosphocytidyl-2C-methyl-D-erythritol synthase; *CMK*: 4-diphosphocytidyl-2C-methyl-D-erythritol kinase; *MECPS*: 2C-methyl-D-erythritol 4-phosphate cytidylyltransferase; HDS: 1-hydroxy-2-methyl-2-(E)-butenyl-4-diphosphate synthase; *HDR*: 1-hydroxy-2-methyl-2-(E)-butenyl-4-diphosphate reductase, MVA pathway genes: *AACT*: acetoacetyl-CoA thiolase; *HMGS*: 3-hydroxy-3-methylglutaryl-CoA synthase; *HMGR*: 3-hydroxy-3-methylglutaryl-CoA reductase; *MVK*: mevalonate kinase; *PMVK*: phosphomevalonate kinase; *MVD*: mevalonate diphosphate decarboxylase; *IPPI*: isopentenyl-diphosphate isomerase, and prenyl transferases (PT) *FPPS*: farnesyl diphosphate synthase; *GPP*S: geranyl diphosphate synthase.

Total mono-*TPS* genes showed similar mean expression pattern trends until the veraison stage among cultivars; however, we observed clear trends of increased expression in Albariño, Gewürztraminer, Riesling, and Viognier and decreased expression in Pinot Noir at harvest compared to veraison (Table S5).

The mean expression of putative mono-*TPS* genes was higher at early developmental stages compared to later stages (Figure 5; Table S5). We observed extensive anthesis-specific mono- *TPS* gene expression (62 genes) across cultivars (Figure 6) including genes from OG1 (putative *(E)-β-*ocimene/myrcene synthases; *VvTPS38*) and OG9 (putative geraniol synthases (*VvTPS52*); Table S4).

**Figure 6:**
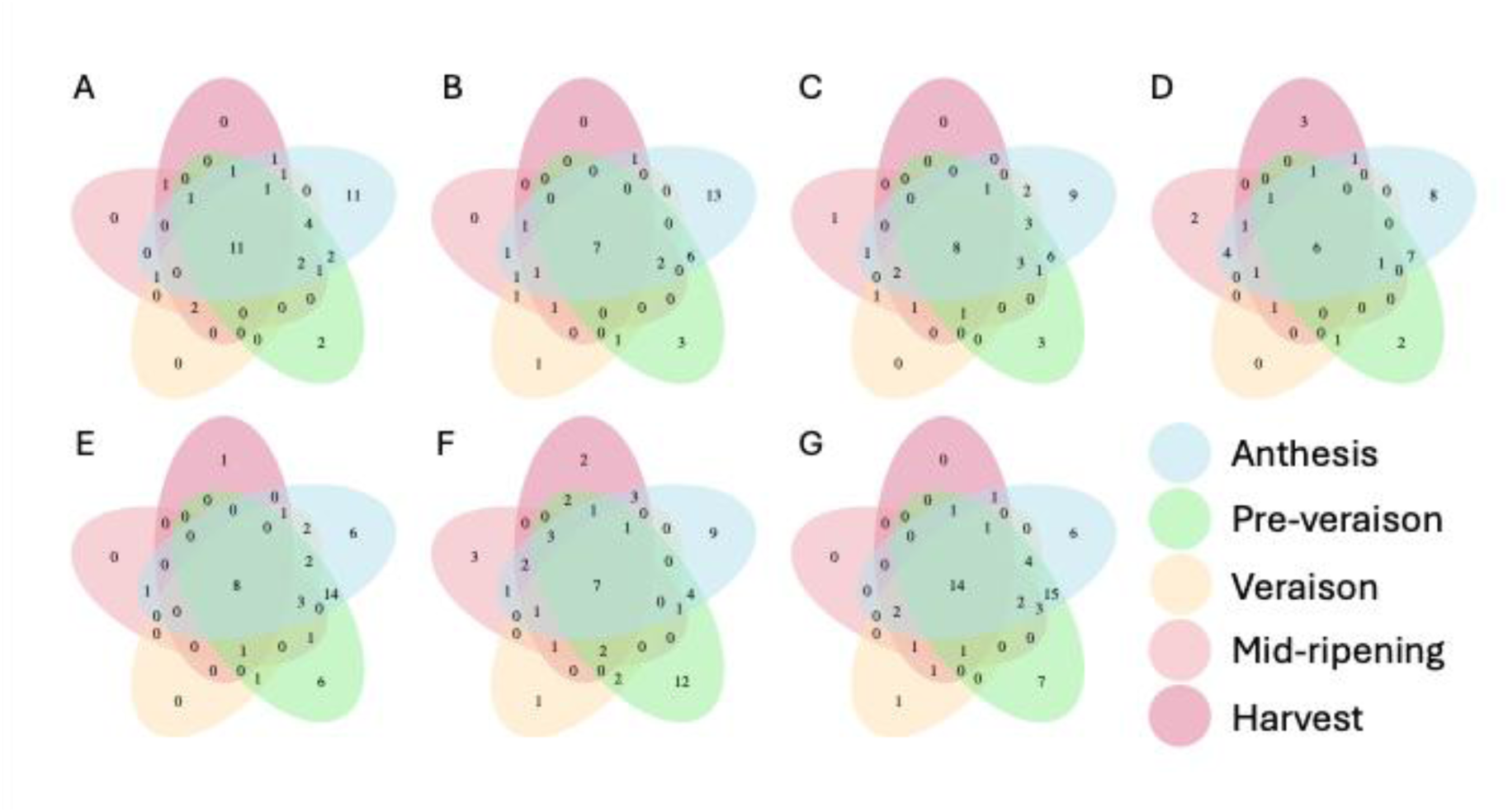
Venn diagrams showing the number of expressed mono-TPS throughout development for (A) Albariño, (B) Cabernet Sauvignon, (C) Fiano, (D) Gewürztraminer, (E) Pinot Noir, (F) Riesling, and (G) Viognier. Colours represent the developmental stage.

Later developmental stages exhibited limited stage-specific expression; however Riesling showed the highest number of mono-*TPS* genes expressed at post-veraison (28 genes), while Cabernet Sauvignon and Pinot Noir exhibited the least (14 and 12 genes, respectively). Similarly, on average Albariño, Fiano, Gewürztraminer, Riesling, and Viognier exhibited 5.1- and 15.16-fold higher expression than Cabernet Sauvignon and Pinot Noir during mid-ripening and harvest, respectively (Table S6). Notably, at harvest Gewürztraminer exhibited 6.15-fold higher mean mono-*TPS* gene expression compared to the average of other cultivars (Figure 5; Table S6).

No expression was observed for private mono-*TPS* genes and dispensable mono-TPS OG15 and OG18 in the flower and berry samples analyzed. Generally, mono-TPS OGs shared temporal expression patterns across cultivars (Figure 5). Exceptions were seen in OG1 (putative *(E)-β*-ocimene/myrcene synthases; *VvTPS38*), in which total Riesling and

Gewürztraminer gene expression increased by 22.95- and 89.38-fold at harvest compared to veraison, respectively while no significant increases were identified for other cultivars (Figure 5; Table S5). Similarly, in OG8 (putative *(E)-β*-ocimene synthases; *VvTPS35*) gene expression increased in Gewürztraminer, Riesling, and Viognier by 137.87-, 34.3- and 3.16-fold at harvest compared to veraison, respectively. In Cabernet Sauvignon and Pinot Noir, OG11 (putative (−)- *α*-terpineol synthases; *VvTPS39*) genes were only expressed before the onset of ripening, whereas Albariño, Fiano, Gewürztraminer, Riesling, and Viognier showed expression of OG11 genes at veraison and post-veraison.

### d. Identification of key terpenoids during berry ripening across cultivars

We identified twenty-eight free terpenoids, including twenty-four monoterpenoids and four sesquiterpenoids in the developing flowers and berries (Table S7, S8). Terpenoid concentrations were dynamic in flowers and berries. Total free terpenoid concentrations were highest at anthesis in all cultivars (ANOVA, p-value<0.05; >928 µg kg^-1^), whereas total bound terpenoids were highest at anthesis for Cabernet Sauvignon, Pinot Noir, and Riesling, at anthesis and harvest for Albariño, Fiano, and Viognier, and at harvest for Gewürztraminer (Table S7). During early berry development the total terpenoid concentrations decreased in all cultivars, after which we observed a tendency of increasing concentrations in ripening berries for Albariño, Fiano, Gewürztraminer, Riesling, and Viognier, and constant or decreasing concentrations for Cabernet Sauvignon and Pinot Noir. Based on the divergent accumulation patterns during berry ripening, cultivars were classified as high-terpenoid (Albariño, Fiano, Gewürztraminer, Riesling, and Viognier), and low-terpenoid (Cabernet Sauvignon and Pinot Noir) cultivars, respectively. Notably, even within the grouping of the high-terpenoid cultivars, we could observe a range in concentrations, with Gewürztraminer accumulating significantly higher concentration of terpenoids in ripening berries compared to other cultivars (Table S8).

The developmental stage was a major driver of differences in terpenoid concentrations. While similar accumulation patterns were detected across cultivars, a distinct divergence occurred post-veraison for selected terpenoids (Figure 7).

**Figure 7:**
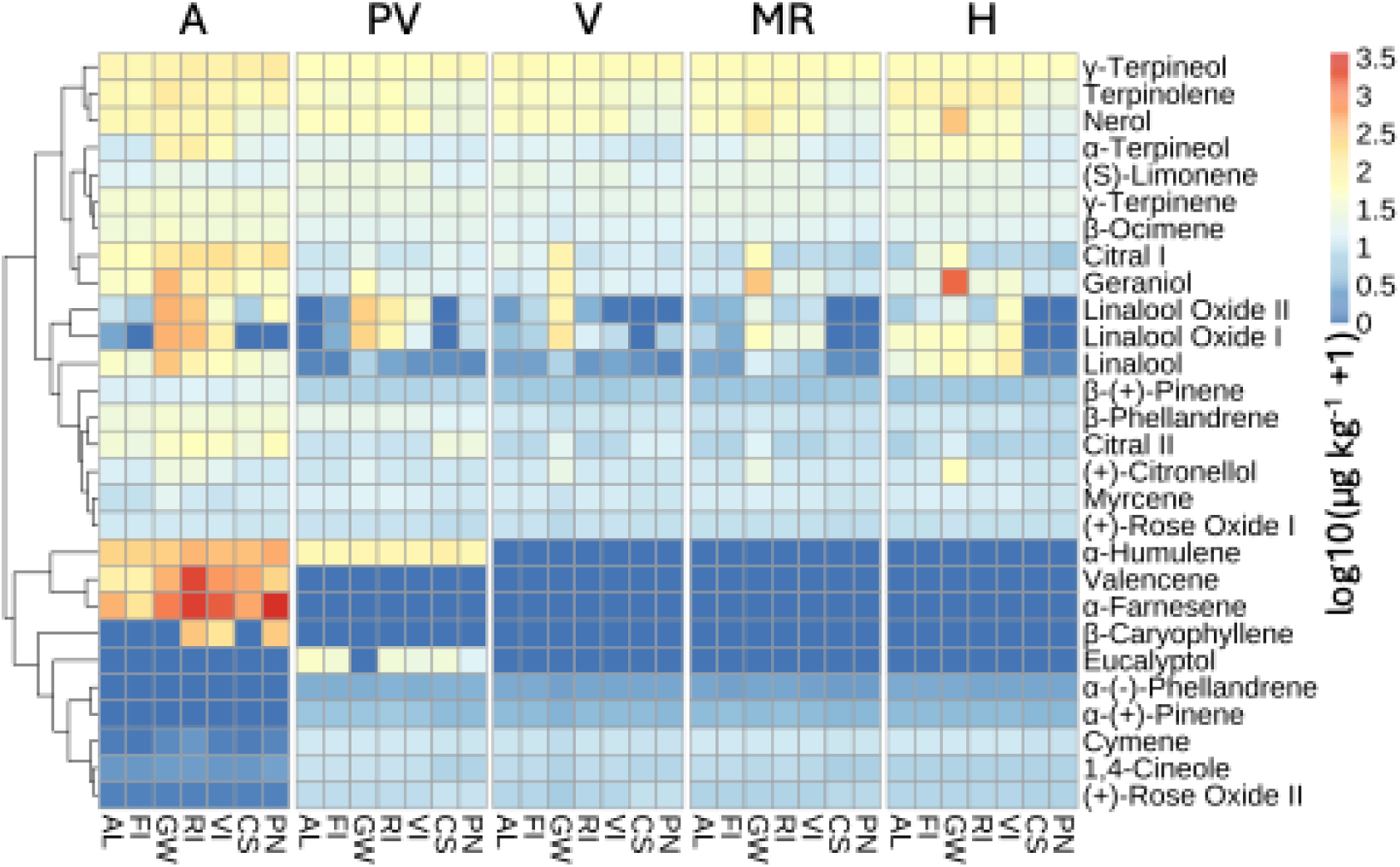
Heatmap of log₁₀-transformed terpenoid concentrations (µg kg^-1^ fresh weight) across different grape varieties. Metabolite concentrations represent the sum of free and bound terpenoids. Concentrations are color-coded, with red indicating high concentrations and blue representing low concentrations. Metabolites with similar accumulation patterns are grouped in the dendrogram. Developmental stages are abbreviated as follows: anthesis (A), pre-veraison (PV), veraison (V), mid-ripening (MR), and harvest (H). The analysed grape varieties include Albariño (AL), Cabernet Sauvignon (CS), Fiano (FI), Gewürztraminer (GW), Pinot Noir (PN), Riesling (RI), and Viognier (VI).

In the developing berries of low-terpenoid cultivars, individual terpenoid concentrations remained stable or decreased during ripening. In contrast, high-terpenoid cultivars exhibited pronounced increases in free and bound terpenoid concentrations at harvest compared to veraison for free *α*-terpineol (6.65-fold mean across cultivars), geraniol (3.14-fold), linalool (8.22-fold), terpinolene (3.75-fold), bound geraniol (13.44-fold), linalool, and linalool oxide I, the last two were below the limit of detection at veraison. Additionally, we identified significant increases in bound (+)-citronellol (3.01-fold) and nerol (13.6-fold) in Gewürztraminer, while free linalool oxide I (3.35-fold) and II (15.97-fold) increased in Albariño and Viognier, and bound linalool oxide II was below the limit of detection at veraison but not at harvest in Viognier. Despite sharing similar accumulation patterns, high-terpenoid cultivars differed significantly at harvest in their concentrations of free geraniol, linalool oxide I, linalool oxide II, 1,4-cineole, y-terpineol, nerol, (+)-rose oxide isomer I and II, and bound *α*-terpineol, geraniol, linalool, linalool oxide I, linalool oxide II, (+)-citronellol, and nerol. Among the cultivars studied, Gewürztraminer was characterized by consistently highest concentrations of both total free and bound terpenoids in the berries, largely driven by geraniol, nerol, (+)-citronellol, and citral isomer I concentrations (Figure 7, Table S8). To identify the key terpenoids driving cultivar-specific ripe berry aroma, we established two inclusion criteria: (1) at least a 2-fold concentration increase from veraison to harvest in at least one cultivar or (2) statistically significant and substantial (>10 µg kg⁻¹) variation in concentration among high-terpenoid cultivars at harvest. Based on these thresholds, our data suggest that free and bound *α*-terpineol, geraniol, linalool, linalool oxide I and II, free terpinolene, and bound (+)-citronellol and nerol are key terpenoids that might impact cultivar-specific aroma profiles at harvest. Notably, all these selected terpenoids were found in significantly higher concentrations in ripe berries of high-terpenoid compared to low-terpenoid cultivars.

### e. Transcript-metabolite and transcript-transcript correlations

To identify putative transcript-metabolite relationships, we conducted a Pearson correlation analysis between metabolite concentrations and terpenoid biosynthesis gene expression from anthesis to harvest across all cultivars. We identified significant correlations between total terpenoid accumulation and the expression of several pathway genes (Table 2).

**Table 2:** Pearson correlation results for relationship of cumulative pathway gene expression and total terpenoid concentrations throughout flower and berry development across varieties.

| Gene | R-value | adjusted p-value |
| --- | --- | --- |
| *DXS* | 0.29 | 0.158 |
| *DXS1* | 0.30 | 0.146 |
| *DXR* | 0.24 | 0.220 |
| *MCT* | 0.46 | **0.015\*** |
| *CMK* | -0.07 | 0.699 |
| *MECPS* | -0.16 | 0.387 |
| *HDS* | -0.55 | **0.002\*** |
| *HDR* | -0.39 | 0.053 |
| *GPPS* | -0.25 | 0.217 |
| *AACT* | -0.17 | 0.387 |
| *HMGS* | 0.37 | 0.062 |
| *HMGR* | 0.55 | **0.002\*** |
| *MVK* | 0.33 | 0.099 |
| *PMVK* | -0.17 | 0.387 |
| *MVD* | 0.75 | **<0.001\*** |
| *IPPI* | -0.08 | 0.671 |
| *FPPS* | 0.58 | 0.002 |
| *TPS* | 0.79 | **<0.001\*** |

Within the MEP pathway, total terpenoids correlated moderately with *MCT* (R = 0.46, p-adjusted = 0.015) and negatively with *HDS* (R = −0.55, p-adjusted = 0.002; Table 2). Within the MVA pathway, significant positive correlations were observed for *HMGR* (R = 0.55, p-adjusted = 0.002), *MVD* (R = 0.75, p-adjusted < 0.001), and *FPPS* (R = 0.58, p-adjusted = 0.001). While *MVD* was the only upstream pathway gene to show a strong correlation (R > 0.70, p-adjusted < 0.05), the strongest positive correlation overall was identified between total *TPS* (mono- and sesqui-*TPS*) expression and total terpenoid concentration (R = 0.79, p-adjusted < 0.001). This suggests that downstream *TPS* expression, rather than upstream pathway regulation, serves as a driver of terpenoid accumulation in flowers and berries. To identify the *TPS* genes strongly associated with the aroma accumulation in the cultivars analyzed, a transcript-metabolite correlation between individual *TPS* and metabolites was conducted.

Strong positive correlations (R ≥ 0.80, p-adjusted < 0.05) between individual *TPS* gene expression and their putative terpenoid products were identified and visualized in a correlation network (Figure 8).

**Figure 8:**
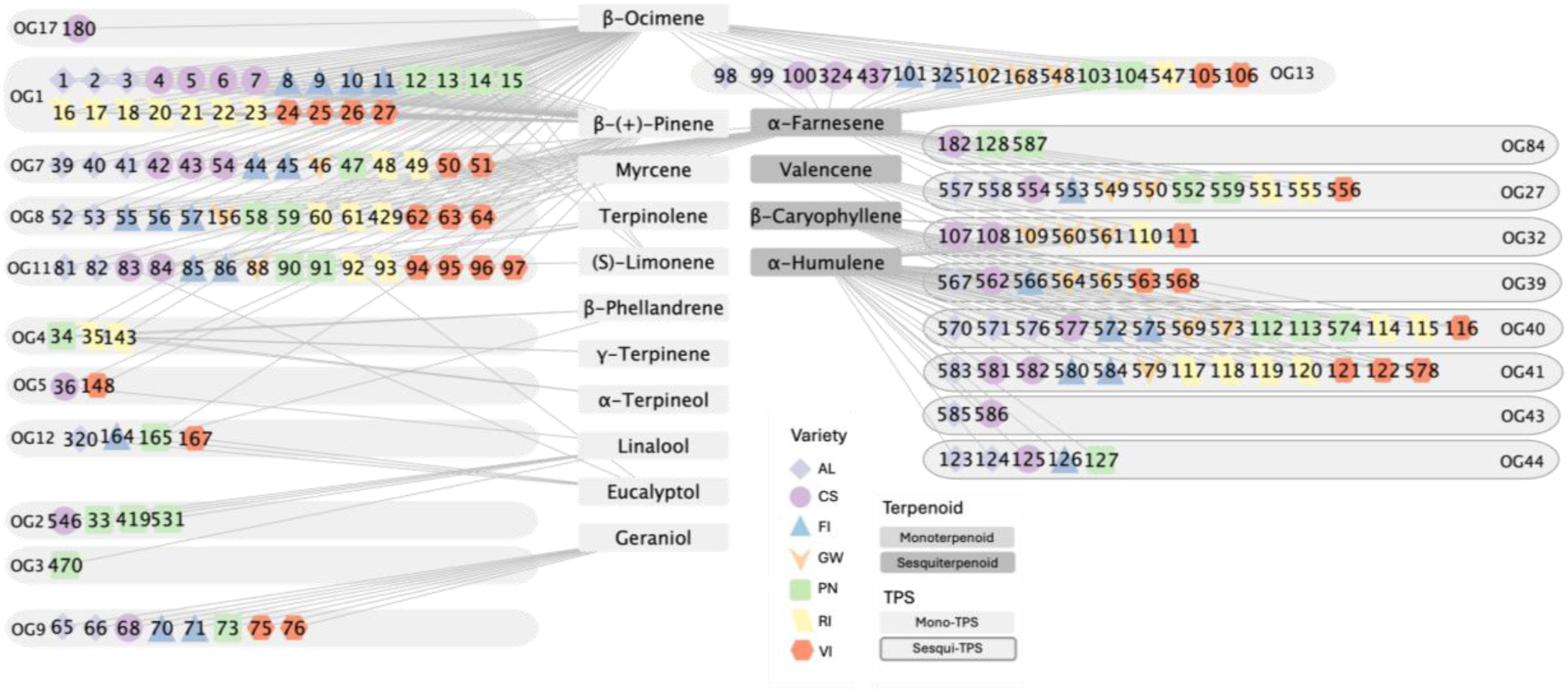
Network representation of transcript-metabolite Pearson correlation between *TPS* genes and associated terpenoids in Albariño (AL), Cabernet Sauvignon (CS), Fiano (FI), Gewürztraminer (GW), Pinot Noir (PN), Riesling (RI), and Viognier (VI). *TPS* genes are colour coded based on varieties and represented as members of orthogroups (OGs). The numbers of individual *TPS* genes can be cross-referenced with column ‘Map_numbers’ in Table S4. Only strong positive correlations (R ≥ 0.8, p-adjusted < 0.05) are shown. Terpenoids are the sum of free and bound terpenoids.

The expression of putative *β*-ocimene synthases (OG1, OG7, and OG8), geraniol synthases (OG9), linalool synthases (OG2 and OG3), *α*-terpineol synthases (OG11), *β*-phellandrene synthases (OG4), *α*-pinene synthases (OG5), *β*-ocimene/*α*-farnesene synthases (OG13), *α*-humulene synthases (OG44), valencene synthases (OG32), *α*-bergamotene synthases (OG27), *β*-caryophyllene synthases (OG39, OG40, OG41, and OG43), and *α*-farnesene synthases (OG84) correlated with their respective product concentrations. However, beyond transcript-associated metabolite correlations, numerous strong positive correlations between transcripts and metabolites that are not their putative products have been identified (Table S9), suggesting co-regulation of terpenoid biosynthesis gene expression. A transcript-transcript Pearson correlation analysis revealed strong positive correlations (R ≥ 0.8, p-adjusted < 0.05) between expression patterns of terpenoid biosynthesis genes (Figure S6). Correlations between MEP and MVA pathway gene expressions were largely between MVA gene *HMGR* and prenyl transferase *FPPS* with early MEP genes *DXS*, *DXR*, and *MCT*. Additionally, many strong positive correlations were identified between MEP and MVA pathway gene expressions with mono-*TPS* gene expressions, proposing high levels of co-regulation between terpenoid biosynthesis gene expression.

To evaluate whether the expanded *TPS* CNV drives cultivar-specific aroma profiles through a gene dosage effect, we analyzed the relationship between CNV, gene expression, and metabolite concentrations across development (Figure S7). Interestingly, no strong positive correlations between *TPS* CNV and their total expression level or final metabolite concentration across developmental stages were identified. Instead, CNV effects were developmental stage specific. At anthesis and pre-veraison, the CNVs of OG11 (putative (-)-*α*-terpineol synthase; *VvTPS39*) and OG3 (putative (*3S*)-linalool/(*E*)-nerolidol synthase; *VvTPS54*) correlated with the accumulation of their respective putative products. Likewise, CNV correlated with gene expression at specific developmental points: OG2 (putative (*3S*)-linalool/(*E*)-nerolidol/(*E,E*)-geranyl linalool synthase; *VvTPS57/58/61/63*) CNV correlated with its expression at anthesis, while OG5 (putative (+)-*α*-pinene synthase, *VvTPS44*) CNV correlated with expression at mid-ripening and harvest. These stage-specific correlations suggest that while genomic structural expansion provides potential for high terpenoid diversity across cultivars, there is a strong developmental control over expression.

### f. Functional characterization of candidate *TPS* genes in *N. benthamiana*

The candidate mono-TPS groups OG12, OG14, OG16, and OG17 lacked previously characterized representative genes but were actively expressed in our dataset (Figure 5; Table S4). To elucidate how their expression contributes to the berry aroma profile, we performed functional characterizations using transient *TPS* expression and metabolite analysis in *N. benthamiana*. For each OG, we selected two representative genes from different cultivars for functional characterization. Additionally, a Gewürztraminer-derived OG8 gene, which has 100% sequence identity with a previously characterized *(E)-β*-ocimene synthase (*VvTPS35*; Martin et al., 2010), was re-characterized. This gene was highly expressed at harvest. It lacked a typical transit peptide characteristic of plastid-localized mono-TPS but instead was predicted to have cytosolic localization (TargetP), which warranted functional validation. As a positive control, we included a *Catharanthus roseus* TPS (*CrGES*) previously characterized as a functional geraniol synthase in *N. benthamiana* (Dudley et al., 2022; Höfer et al., 2013). To maximize upstream precursor availability, candidate *TPS* genes were co-expressed with rate-limiting enzymes of the terpenoid biosynthetic pathways, including DXS (*CLA1*; *A. thaliana*) for the MEP pathway and HMGR (*HMG1*; *A. thaliana*) for the MVA pathway, as well as GPPS (*PaGPPS*; *Picea abies*). The *p19* silencing suppressor was included in all transformations, including the controls.

Expression of OG14 (*VITVvi_vGewFPS47_v1.0.hap2.chr13.ver1.0.g484560, VITVvi_vViogFPS05_v1.0.Hap2.chr13.ver1.0.g472970*) and OG16 (*VITVvi_vCabSauv08_v2.0.hapCF.chr13.ver2.0.g196620, VITVvi_vGewFPS47_v1.0.hap1.chr13.ver1.0.g193050*) *TPS* candidate genes in *N. benthamiana* did not result in detectable terpenoid formation compared to controls.

Expression of Fiano OG12 gene *VITVvi_vFianoFPS02.1_v1.0.Hap1.chr13.ver1.0.g193300* resulted in terpenoid formation that differed from the controls (Figure 9B-C) and with product profiles that were affected by co-expression of precursor pathway genes (Table S10).

**Figure 9:**
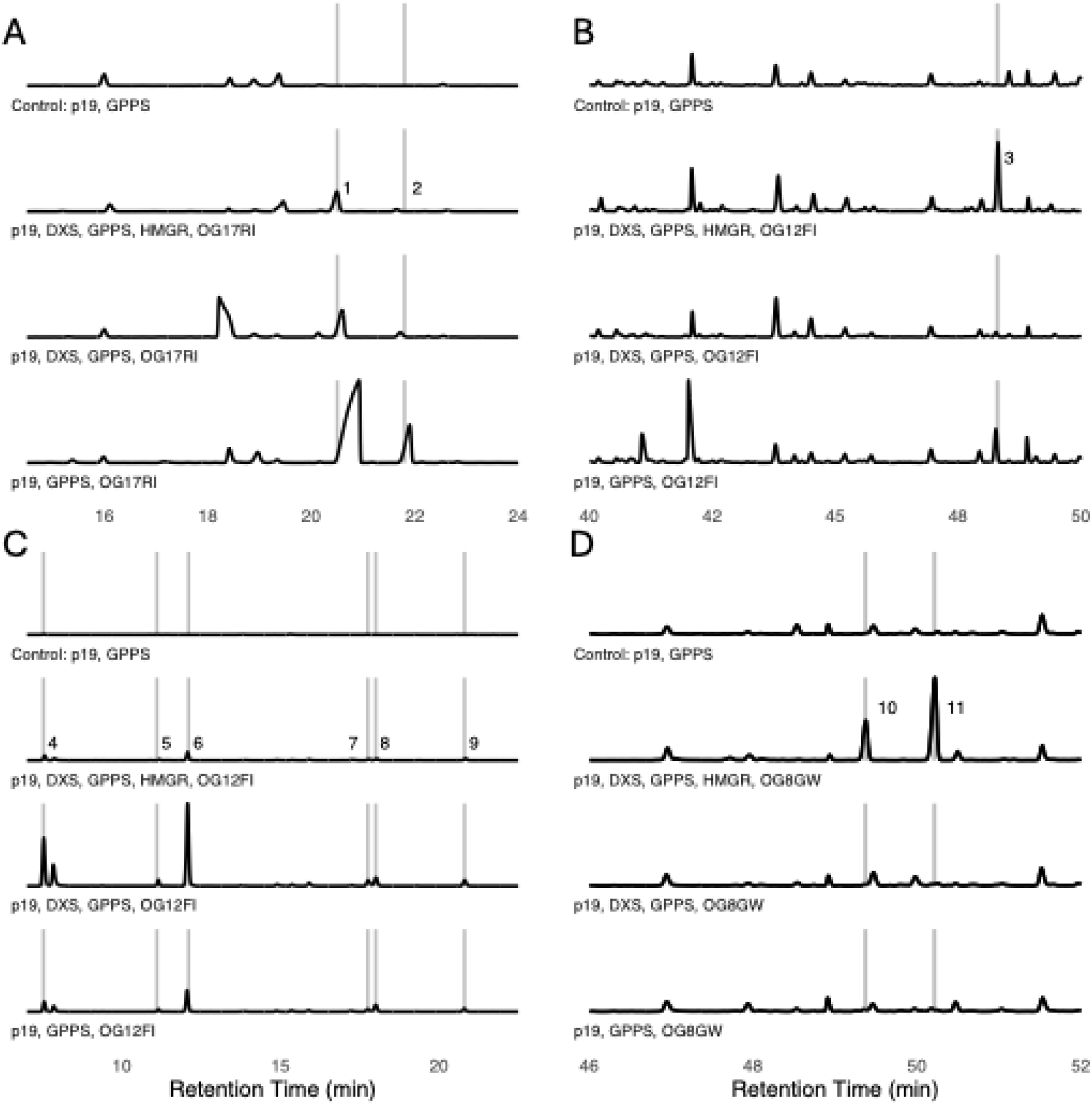
Chromatograms of secondary metabolite products from *Nicotiana benthamiana* leaves infiltrated with the p19 suppression of silencing vector, *GPPS* (geranyl diphosphate synthase), *DXS* (1-deoxy-D-xylulose 5-phosphate synthase) and *HMGR* (3-hydroxy-3-methylglutaryl-CoA reductase) together with (A) TIC of OG17 Riesling gene (*VITVvi_vRiesFPS24_v1.1.hap2.chr12.ver1.0.g502900*), (B) TIC of OG12 Fiano gene (*VITVvi_vFianoFPS02.1_v1.0.Hap1.chr13.ver1.0.g193300*), (C) EIC93 of OG12 Fiano gene (*VITVvi_vFianoFPS02.1_v1.0.Hap1.chr13.ver1.0.g193300*), (D) OG8 Gewürztraminer gene (*VITVvi_vGewFPS47_v1.0.hap1.chr12.ver1.0.g172600*). Peaks different to control were 1, *trans-β*-ocimene; 2, *cis-β*-ocimene; 3, *α*-terpineol; 4, *α*-pinene; 5, *β*-pinene; 6, sabinen; 7, *β*-phellandrene; 8, eucalyptol; 9, *γ*-terpinene; 10, *(Z, E)-α*-farnesene; 11, *(E,E)-α*-farnesene.

When *VITVvi_vFianoFPS02.1_v1.0.Hap1.chr13.ver1.0.g193300* was co-expressed with *DXS*, *GPPS*, and *HMGR*, the terpenoid profile was dominated by *α*-terpineol (57.2 ± 21.42% total peak area; mean ± SE) and cymene (20.27 ± 11.34%), and minor peaks (<10%) of *γ*-terpinene, *α*-pinene, eucalyptol, sabinene, *β*-phellandrene, and *β*-pinene. However, when the same gene was co-expressed with only with *DXS* and *GPPS*, the profile changed to predominantly cymene (34.15 ± 7.27% peak area), *α*-pinene (21.3 ± 4.67%), sabinene (11.82 ± 5.07%), and eucalyptol (11.14 ± 2.04%), while *α*-terpineol, *γ*-terpinene, *β*-phellandrene, and *β*-pinene were detected as minor products. Finally, when this gene was co-expressed only with *GPPS*, the product profile was dominated by *α*-terpineol (20.78 ± 13.77% peak area), *α*-pinene (18.84 ± 7.16%), sabinene (16.78 ± 6.12%), eucalyptol (16.7 ± 0.32%), cymene (15.0 ± 1.09%), and minor peaks of *γ*-terpinene and *β*-pinene. In summary, under all three conditions the same terpenoids were produced in *N. benthamiana* expressing Fiano OG12 gene *VITVvi_vFianoFPS02.1_v1.0.Hap1.chr13.ver1.0.g193300*, however the relative proportions of individual products shift based on upstream pathway gene expression. Under identical experimental conditions, the Riesling OG12 ortholog *VITVvi_vRiesFPS24_v1.1.hap1.chr13.ver1.0.g211380*, failed to produce detectable terpenoid levels compared to the control. Notably, this Riesling gene shared 100% amino acid sequence identity with Albariño (*VITVvi_vAlbaFPS03.1_v1.0.Hap2.chr13.ver1.0.g471560*), Cabernet Sauvignon (*VITVvi_vCabSauv08_v2.0.hapSB.chr13.ver2.0.g491410*), Gewürztraminer (*VITVvi_vGewFPS47_v1.0.hap1.chr13.ver1.0.g193070*), Pinot Noir (*VITVvi_vPinNoir123_v1.0.hap1.chr13.ver1.0.g197740*) and Viognier (*VITVvi_vViogFPS05_v1.0.Hap1.chr13.ver1.0.g188770*) genes within OG12.

OG17 Riesling gene *VITVvi_vRiesFPS24_v1.1.hap2.chr12.ver1.0.g502900*, which shared 100% amino acid sequence identity with Cabernet Sauvignon (*VITVvi_vCabSauv08_v2.0.hapCF.chr12.ver2.0.g174790*), produced similar proportions of *β*-ocimene isomers (Figure 9A; Table S10), averaging 90.1 ± 2.35% *trans-β*-ocimene and 9.90 ± 2.35% *cis-β*-ocimene, across all three pathway co-expression combinations, whereas OG17 Viognier gene *VITVvi_vViogFPS05_v1.0.Hap1.chr12.ver1.0.g168900* showed no detectable terpenoid production compared to the control.

Expression of Gewürztraminer OG8 gene *VITVvi_vGewFPS47_v1.0.hap1.chr12.ver1.0.g172600*, which has 100% sequence identity with previously characterized (*E*)-*β*-ocimene synthase (*VvTPS35*; Martin et al., 2010), in the *N. benthamiana* system revealed exclusive production of sesquiterpenoids (*Z, E*)*-α*-farnesene and (*E,E*)-*α*-farnesene isomers in all three co-expression assays described above (Figure 9D). Notably, the total peak area of these sesquiterpenoids was over 15-fold larger when *VITVvi_vGewFPS47_v1.0.hap1.chr12.ver1.0.g172600* was co-expressed with the MVA pathway rate-limiting gene *HMGR* compared to co-expression with the plastidial MEP pathway gene *DXS* and *GPPS* alone.

## 5. Discussion

Eight monoterpenoids were identified as major contributors to terpenoid profile differences among the berries of seven major grapevine cultivars. During berry development, total monoterpenoid concentrations decreased until the onset of ripening (veraison). Post-veraison, concentrations remained low or continued to decline in low-terpenoid cultivars, whereas they increased significantly in high-terpenoid cultivars. This divergent accumulation pattern aligns with the historical categorization of Cabernet Sauvignon and Pinot Noir as neutral or non-aromatic cultivars, and Albariño, Fiano, Gewürztraminer, Riesling, and Viognier as aromatic cultivars (Kalua and Boss, 2009; Matarese et al., 2013; Mateo and Jiménez, 2000). Our results suggest that the concentrations of free and bound *α-*terpineol, geraniol, linalool, linalool oxide I and II, free terpinolene, and bound (+)-citronellol and nerol serve as the key contributors of aroma divergence among the ripe berries of these aromatic cultivars. The importance of specific subsets of these volatile compounds in aromatic grape fragrance has been noted previously for individual cultivars (Bosman et al., 2023; Girard et al., 2002; Kovalenko et al., 2021; Lin et al., 2019; X. Liu et al., 2022; Marais, 1983). In particular, it has been shown that the characteristic citrus and floral notes of Riesling and the stone fruit like aroma of Viognier are largely determined by linalool, geraniol, and nerol concentrations (Marais, 1983; Siebert et al., 2018; Wang et al., 2019), whereas the floral and lychee aroma of Gewürztraminer is produced by geraniol, α-terpineol, (+)-citronellol, and (+)-rose oxide levels (Kovalenko et al., 2021; Martin et al., 2012). Our study extends the list of key compounds for the seven cultivars considered and suggest that their differences in concentrations could determine the difference in varietal aromas of berries and wines.

DXS activity represents a rate-limiting step in terpenoid accumulation in plants (Tholl, 2015), and the Muscat *DXS1* gain-of-function mutation has been identified as a key driver of terpenoid accumulation (Battilana et al., 2011; Emanuelli et al., 2010). This mutation increases precursor availability within the MEP pathway enabling elevated terpenoid accumulation in Muscat cultivars. However, none of the *DXS1* genes in the seven cultivars analyzed in the present study possess this specific mutation; similarly, numerous other aromatic, non-Muscat cultivars also lack this *DXS1* polymorphism (Emanuelli et al., 2010). Furthermore, neither *DXS1* nor total *DXS* expression correlated with total terpenoid concentrations in our dataset (Figure S7). These findings imply that alternative regulatory mechanisms contribute as drivers to terpenoid accumulation, and that the elevated terpenoid profiles of the high-terpenoid cultivars studied here are independent of the previously characterized *DXS1* mutation.

Our analysis of haplotype-resolved genome assemblies revealed substantial and variable expansions of the *TPS* gene family in different grapevine cultivars, ranging from 53-70 *TPS* genes in Fiano to 86-89 *TPS* genes in Riesling depending on individual haplotypes. Expansions of *TPS* gene families have been reported for genomes of other fruit crop such as strawberry (*Fragaria × ananassa*; Madera et al., 2026), with 75 full length *TPS*, and apple (*Malus domestica*) with 55 putative *TPS* (Nieuwenhuizen et al., 2013), as well as other plants with large and diverse terpenoid profiles like eucalyptus (*Eucalyptus grandis*) with 113 putative *TPS* (Külheim et al., 2015), compared to the relatively smaller *TPS* gene family of the model system *A. thaliana* with 32 *TPS* (Aubourg et al., 2002). The stark variation in grapevine *TPS* copy numbers observed across haplotypes and cultivars in this study highlights the importance of haplotype-resolved reference assemblies to decipher these duplicated regions of highly heterozygous genomes involved in cultivar-specific aroma profiles.

Unexpectedly, cultivar differences in terpenoid accumulation were explained by *TPS* expression rather than by copy number, with gene dosage largely decoupled from both transcript abundance and metabolite levels. While previous literature has linked higher copy numbers to elevated terpenoid accumulation through a dosage effect or hyper-functional TPSs, like the relationship seen between *TPS39* and cyclic monoterpenoids (*α*-terpinene, p-cymene, 1,8-cineole, *γ*-terpinene, *α*-terpinolene, terpinene-4-ol, *α*-terpineol; Bosman et al., 2023), our results suggest that *TPS* CNV is not strongly associated with strength of expression or terpenoid accumulation throughout development (Figure S7). Instead, our findings suggest that the expression of these expanded *TPS* gene sets is under tight developmental control. Indeed, the total expression of *TPS* correlated strongly with the total terpenoid accumulation throughout berry development, suggesting a key role in shaping terpenoid-based aroma profiles. Previous QTL mapping in cross-populations identified loci containing *TPS* clusters as determinants of terpenoid traits (Bosman et al., 2023; Emanuelli et al., 2010). Recently, Lin et al. (2026) identified an aroma-associated QTL harboring (3S)-linalool/nerolidol synthases (*VvTPS54*) and demonstrated that their expression profile was strongly associated with linalool accumulation in grape berries at harvest. An additional, non-exclusive possibility is that duplicated *TPS* copies within orthogroups have undergone sub- or neofunctionalization (Chen et al., 2011; Force et al., 1999), so that copy number does not equate to functional redundancy. Testing functional divergence among paralogues, rather than the orthologous copies compared here, was beyond the scope of this study but would clarify how the expanded family maps onto terpenoid output.

Within each OG, the timing of *TPS* expression was conserved across cultivars, whereas different OGs exhibited distinct temporal expression patterns. Interestingly, we observed extensive anthesis-specific mono-TPS expression; however, the biological role of terpenoids in domesticated grapevine flowers remains unknown (Smit et al., 2019). Deviations from the shared expression patterns and differences in strength of expression within an OG are likely contributing to the different volatile concentrations that we observed among cultivars. For instance, OG11 (putative *α*-terpineol synthase; *VvTPS39*) was expressed at or post-veraison in high-terpenoid cultivars Albariño, Fiano, Gewürztraminer, Riesling, and Viognier, whereas we only observed expression at anthesis and pre-veraison for low-terpenoid cultivars Cabernet Sauvignon and Pinot Noir. Matching this expression pattern, we found on average 16.29-fold higher *α*-terpineol concentrations – the primary product of OG11 *TPS* – at harvest in high-terpenoid cultivars compared to low-terpenoid cultivars. In-depth knowledge of these terpenoid biosynthesis gene expression patterns can identify genes critical for terpenoid accumulation, aiding breeding efforts that target aroma improvements.

Individual transcript-metabolite correlation analysis supported the previously reported *in vitro* characterization of *TPS* genes, whose putative functions were inferred from characterized genes in OGs, through strong alignment with corresponding terpenoid concentrations (Figure 8). Notably, expression levels of specific putative *β*-ocimene synthases (OG7), *α*-terpineol synthases (OG11), *β*-ocimene/*α*-farnesene synthases (OG13), *α*-bergamotene synthases (OG27), and *β*-caryophyllene synthases (OG40) correlated with associated metabolites across all cultivars throughout development (Figure 8). This further supports that the expression level of *TPS* genes is strongly associated with terpenoid accumulation. However, we also encountered widespread, strong correlations between specific *TPS* transcripts and terpenoids that do not align with their predicted enzymatic products (Table S9). A subsequent transcript-transcript network correlation analysis revealed extensive co-regulation among independent terpenoid biosynthesis genes (Figure S6). This includes correlations between MEP/MVA pathway genes, prenyl transferases and *TPS* genes, suggesting that pathway, prenyl transferase, and *TPS* genes are regulated in a similar way. Interestingly, the expression of the *MYB24* transcription factor – previously identified as a *TPS* regulator (Chen et al., 2021; Savoi et al., 2016) – correlated with the expression of terpenoid biosynthesis genes in all cultivars apart from Viognier (Figure S6), proposing a potential conserved regulatory role across cultivars. This extensive co-regulation observed among terpenoid biosynthesis genes limits the ability to infer *TPS* function solely from transcript–metabolite correlations.

To overcome this limitation, two expressed *TPS* candidate genes from each OG without putative functional designation (OG12, OG14, OG16, and OG17) were selected for heterologous expression in *N. benthamiana*. Plant TPS enzymes frequently exhibit substrate plasticity, with product profiles shifting depending on substrate availability (Martin et al., 2010; Martin and Bohlmann, 2004; Matarese et al., 2013). To maximize substrate availability for mono-TPS and sesqui-TPS, we co-overexpressed rate-limiting upstream pathway genes of the MVA pathway (*HMGR*), the MEP pathway (*DXS*) and the potentially rate-limiting prenyl transferase *GPPS*. For the OG14, OG16, OG12 *VITVvi_vRiesFPS24_v1.1.hap1.chr13.ver1.0.g211380*, and OG17 *VITVvi_vRiesFPS24_v1.1.hap2.chr12.ver1.0.g502900* candidate genes, no detectable terpenoids were produced in *N. benthamiana*. Expression of Fiano OG12 gene *VITVvi_vFianoFPS02.1_v1.0.Hap1.chr13.ver1.0.g193300* yielded a mixture of *α*-terpineol, cymene, *α*-pinene, *β*-pinene, sabinene, *β*-phellandrene, eucalyptol, and *γ*-terpinene (Figure 9B-C), confirming that members of the TPS-b subfamily often function as multi-product mono-TPS enzymes with GPP as the substrate (Martin et al., 2010; Martin and Bohlmann, 2004). Riesling OG17 gene *VITVvi_vRiesFPS24_v1.1.hap2.chr12.ver1.0.g502900* produced *trans-β-* ocimene as its primary catalytic product. Additionally, we functionally evaluated the Gewürztraminer gene *VITVvi_vGewFPS47_v1.0.hap1.chr12.ver1.0.g172600*, which shares 100% amino acid sequence identity with previously characterized *(E)-β-*ocimene synthase (*VvTPS35*; Martin et al., 2010), but lacked a full-length transit peptide. Co-expression with upstream MEP and MVA pathway genes (*DXS* and *HMGR*) *plus GPPS* resulted in a large, 15- fold increase in total *α*-farnesene production compared to co-expression with only the *GPPS* and *DXS* genes (Figure 9D). Because *α*-farnesene belongs to the group of sesquiterpenes, which are typically synthesized from precursors of the MVA pathway in the cytosol, the drastic increase in yield upon *HMGR* overexpression strongly indicates that this enzyme accesses the cytosolic component of the terpenoid biosynthesis *in vivo*. This implies that the enzyme is retained within the cytoplasm, likely due to a shortened and non-functional transit peptide, as predicted by TargetP (Table S4). These findings suggest that when the *TPS* was co-expressed only with *GPPS* and MEP pathway gene *DXS*, *α*-farnesene production relied on the endogenous cytosolic FPP pool, whereas *HMGR* overexpression increased precursor availability and consequently led to substantially higher *α*-farnesene accumulation. Indeed, in this study, the expression of *VITVvi_vGewFPS47_v1.0.hap1.chr12.ver1.0.g172600* correlated strongly (R = 0.83, p-adjusted < 0.001) with the concentration of *α*-farnesene in Gewürztraminer flowers and berries throughout development. Similar spatial separation mechanisms of TPS and their substrates, leading to different terpenoid products, have been reported in other plant species. In *A. thaliana*, AtTPS02 contains a complete transit peptide and therefore is localized to plastids, where it produces *trans-β-*ocimene. Whereas nearly identical but transit peptide-lacking AtTPS03 remains in the cytoplasm and synthesizes *(E,E)-α*-farnesene (Huang et al., 2010). Likewise, a snapdragon linalool synthase lacking a complete transit peptide localized to the cytosol and produced the sesquiterpene nerolidol (Nagegowda et al., 2008). In both studies, the enzymes produced both mono- and sesquiterpenoids *in vitro*. Our results suggest that the grapevine mono-*TPS VITVvi_vGewFPS47_v1.0.hap1.chr12.ver1.0.g172600* follows a similar pattern and produces *α*-farnesene *in vivo*, while the identical *VvTPS35* was characterized as a *β-*ocimene synthase *in vitro* (Martin et al., 2010).

However, predicting subcellular localization remains challenging because transit peptides exhibit substantial sequence variation and length differences, even within the same species (Martin et al., 2010). For instance, in the present study, TargetP predicted plastid targeting for only 43.27% of mono-*TPS* genes belonging to the TPS-b and TPS-g subfamilies, highlighting the need for future *in vivo* localization studies to complement the functional validation of grapevine enzymes. Beyond the individual genes characterized here, the manually curated annotation of 85 *TPS* OGs across seven cultivars provides a phylogenetic framework for assigning putative function to *TPS* genes and prioritizing candidates for validation, though as shown here, orthogroup membership alone does not guarantee conserved catalytic function. Consequently, our findings underscore the importance of haplotype-resolved genome assemblies to accurately discover and functionally characterize novel *TPS* genes.

In summary, this study demonstrates the value of cultivar-specific, haplotype-resolved reference genomes for untangling the complex genomic variation that shapes grape berry aroma profiles. We identified eight key terpenoids driving phenotypic aroma divergence across cultivars and demonstrated that differential *TPS* expression, rather than upstream pathway flux, is strongly associated with cultivar-specific aroma accumulation.

## 6. Conclusion

This comprehensive genomic, transcriptomic, and metabolomic study elucidates the regulation of terpenoid biosynthesis in wine grapes, a key biological process determining fruit quality in one of the world’s most economically important crops. By utilizing cultivar-specific, phased diploid genome assemblies, we performed an in-depth analysis of highly heterozygous genomic regions, which are underrepresented in the inbred near-homozygous reference genome PN40024. This approach revealed substantial terpenoid synthase (TPS) gene copy number variation both between haplotypes (up to 24 gene copies within a cultivar) and among cultivars (ranging from 53 TPS genes in Fiano haplotype II to 89 in Riesling haplotype II), advancing our understanding of the genetic architecture underlying grape aroma. The expression of these *TPS* was identified to be strongly associated with terpenoid accumulation patterns, clearly segregating high-terpenoid-producing cultivars (Albariño, Fiano, Gewürztraminer, Riesling, and Viognier) from low-terpenoid-producing cultivars (Cabernet Sauvignon and Pinot Noir), with this metabolic divergence emerging distinctly during berry ripening rather than at flowering or early developmental stages. We further identified eight terpenoids as key drivers of aroma divergence in ripe berries between cultivars. *In vivo* functional characterization of a novel *β*-ocimene synthase, characterized *α*-farnesene synthase, and multi-product *α*-terpineol synthase in *N. benthamiana* provides deeper insights into the complexity of grape berry aroma and establishes a solid foundation for further functional validation of *TPS* genes in wine grapes.

## Acknowledgments

We thank the UC Davis Genome Center DNA Technologies Core Facility for their assistance with sequencing; Dr Mélanie Massonnet and Dr Mirella Zaccheo for their help with sample processing; Guillermo Garcia Zamora for vineyard management; and Zéphir Gondon for his support in data analysis. We want to acknowledge that generative AI (ChatGPT and Gemini) were used to verify and improve the grammar and clarity of sentences and paragraphs.

## 7. Author contribution

SDC, DC, and MPe: conceptualization; SDC and DC: supervision; MPe, MPa, and SDC: writing; SDC and DC: funding; MPe: sampling; MPe and SKD: data analysis; MPe: data visualization; MPe, AM, NC, MPa: bioinformatic analysis; RF-B: library preparation; MPe, LHK, JB: functional gene characterization. All authors have reviewed and approved the manuscript.

## 10. Data availability statement

Raw RNA-Seq data are available on NCBI SRA under BioProject PRJNA1504372. The phased diploid genome assemblies of Albariño, Fiano, Gewürztraminer, and Viognier and their structural annotations are deposited on Zenodo (DOI: <u>10.5281/zenodo.21418196</u>). Dedicated genome browsers are available at www.grapegenomics.com. Mass spectrometry data are available through MetaboLights (raw data; https://www.ebi.ac.uk/metabolights; study identifier MTBLS15219).

---

**Figure S1:** Basic berry parameters displayed as (A) Total soluble solids, (B) pH, and (C) titratable acidity throughout berry development for seven *V. vinifera* varieties. The varieties are Albariño (AL), Cabernet Sauvignon (CS), Fiano (FI), Gewürztraminer (GW), Pinot Noir (PN), Riesling (RI), and Viognier (VI). The developmental stages are pre-veraison (PV), veraison (V), mid-ripening (MR), and harvest (H). Asterisks indicate significant p-value (p<0.05) testing for differences in measurements between varieties within each developmental stage (ANOVA).

**Figure S2:** Phylogenetic tree (protein sequence) of *TPS* genes of subfamily TPS-a from Albariño, Cabernet Sauvignon, Fiano, Gewürztraminer, Pinot Noir, Riesling, Viognier, PN40024v1, PN40024v4.3, and characterized genes. Nodes are annotated with 1000 bootstrap values.

**Figure S3:** Phylogenetic tree (protein sequence) of *TPS* genes of subfamily TPS-c from Albariño, Cabernet Sauvignon, Fiano, Gewürztraminer, Pinot Noir, Riesling, Viognier, PN40024v1, and PN40024v4.3. Nodes are annotated with 1000 bootstrap values.

**Figure S4:** Phylogenetic tree (protein sequence) of *TPS* genes of subfamily TPS-e from Albariño, Cabernet Sauvignon, Fiano, Gewürztraminer, Pinot Noir, Riesling, Viognier, PN40024v1, and PN40024v4.3. Nodes are annotated with 1000 bootstrap values.

**Figure S5:** Phylogenetic tree (protein sequence) of *TPS* genes of subfamily TPS-f from Albariño, Cabernet Sauvignon, Fiano, Gewürztraminer, Pinot Noir, Riesling, Viognier, PN40024v1, PN40024v4.3, and characterised genes. Nodes are annotated with 1000 bootstrap values.

**Figure S6:** Network representation of transcript-transcript Pearson correlation between terpenoid biosynthesis gene expression patterns for Albariño (AL), Cabernet Sauvignon (CS), Fiano (FI), Gewürztraminer (GW), Pinot Noir (PN), Riesling (RI), and Viognier (VI). Only strong positive correlations (R ≥ 0.8, p-adjusted < 0.05) are shown. *TPS* genes are represented as members of orthogroups (OG). Gene names are abbreviated functions of MEP pathway genes: *DXS*: 1-deoxy-dxylulose-5-phosphate synthase; *DXR*: 1-deoxy-D-xylulose-5-phosphate reductoisomerase; *MCT*: 4-diphosphocytidyl-2C-methyl-D-erythritol synthase; *CMK*: 4-diphosphocytidyl-2C-methyl-D-erythritol kinase; *MECPS*: 2C-methyl-D-erythritol 4-phosphate cytidylyltransferase; HDS: 1-hydroxy-2-methyl-2-(E)-butenyl-4-diphosphate synthase; *HDR*: 1-hydroxy-2-methyl-2-(E)-butenyl-4-diphosphate reductase, MVA pathway genes: *AACT*: acetoacetyl-CoA thiolase; *HMGS*: 3-hydroxy-3-methylglutaryl-CoA synthase; *HMGR*: 3-hydroxy-3-methylglutaryl-CoA reductase; *MVK*: mevalonate kinase; *PMVK*: phosphomevalonate kinase; *MVD*: mevalonate diphosphate decarboxylase; *IPPI*: isopentenyl-diphosphate isomerase, and prenyl transferases *FPPS*: farnesyl diphosphate synthase; *GPP*S: geranyl diphosphate synthase. The gene numbers of all genes can be cross-referenced with column ‘Map_numbers’ in Table S4.

**Figure S7:** Spearman correlation analysis of *TPS* gene copy number variation within OGs with (A) mean *TPS* expression summed for OGs and (B) the mean putative product metabolite concentrations throughout berry development. Developmental stages are abbreviated as anthesis (A), pre-veraison (PV), veraison (V), mid-ripening (MR), and harvest (H). Strength of correlation in indicated by colour scale and Spearman rho value. Asterisk marks significant correlations (p-adjusted < 0.05).

