## Supplementary figures and images for "TPS Expression, Not Copy Number, Explains Aroma Diversity among Grapevine (*Vitis vinifera* L.) Cultivars"

### Figure S1

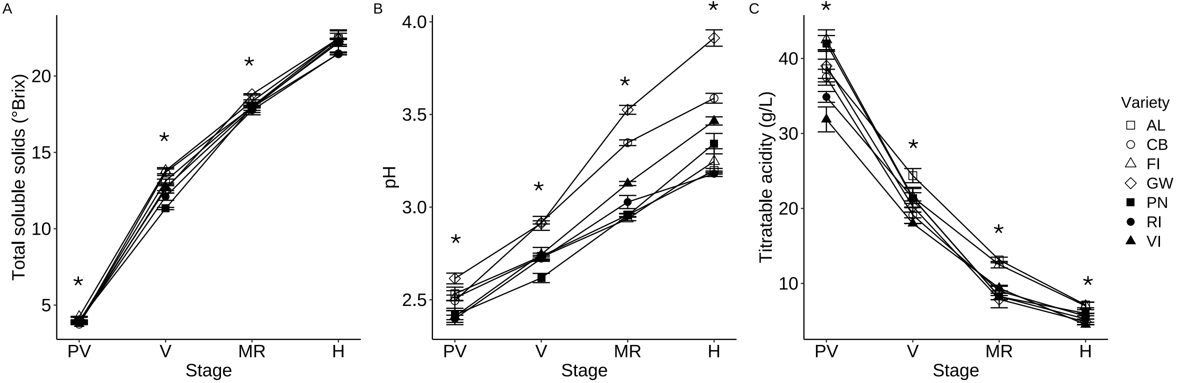

### Figure S2

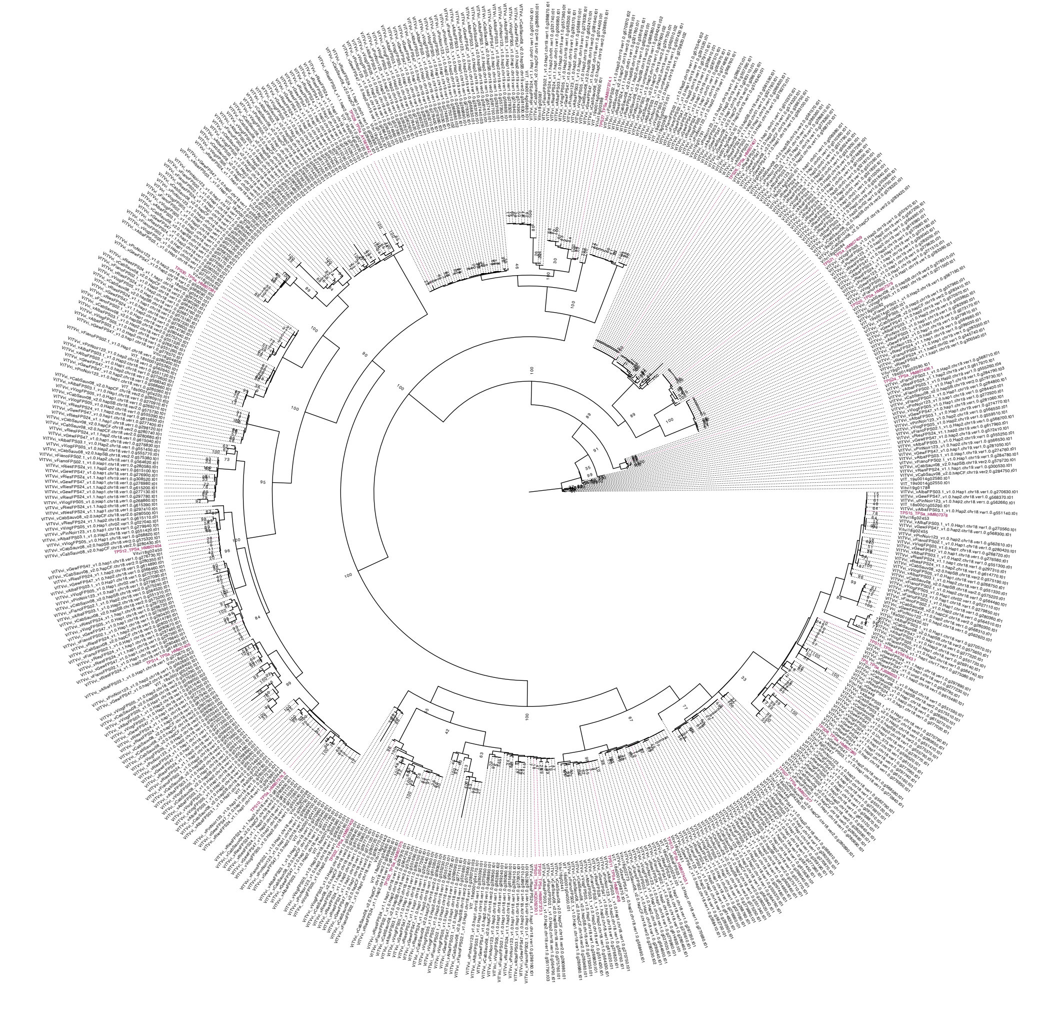

### Figure S3

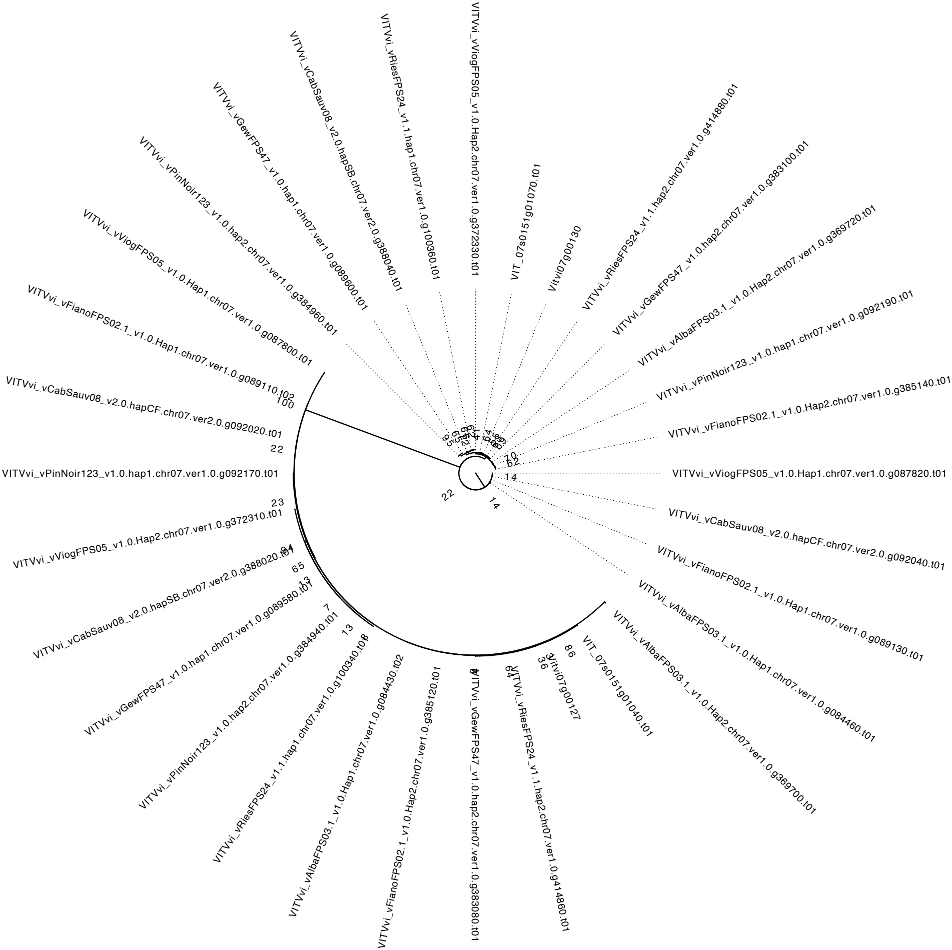

### Figure S4

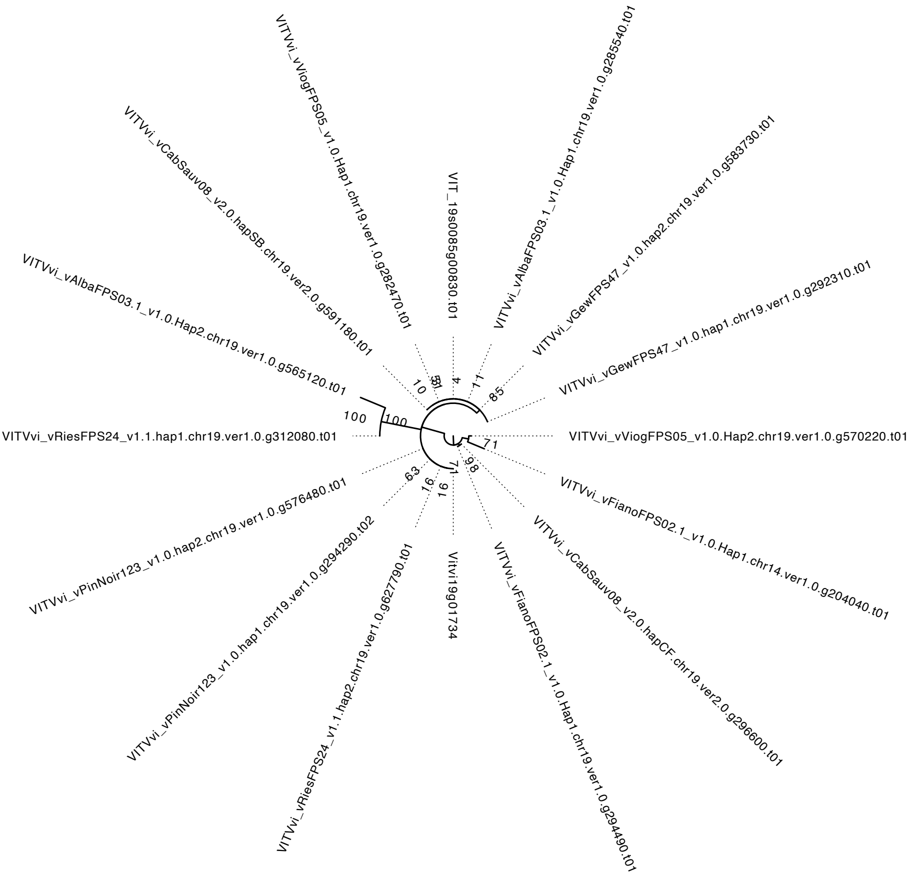

### Figure S5

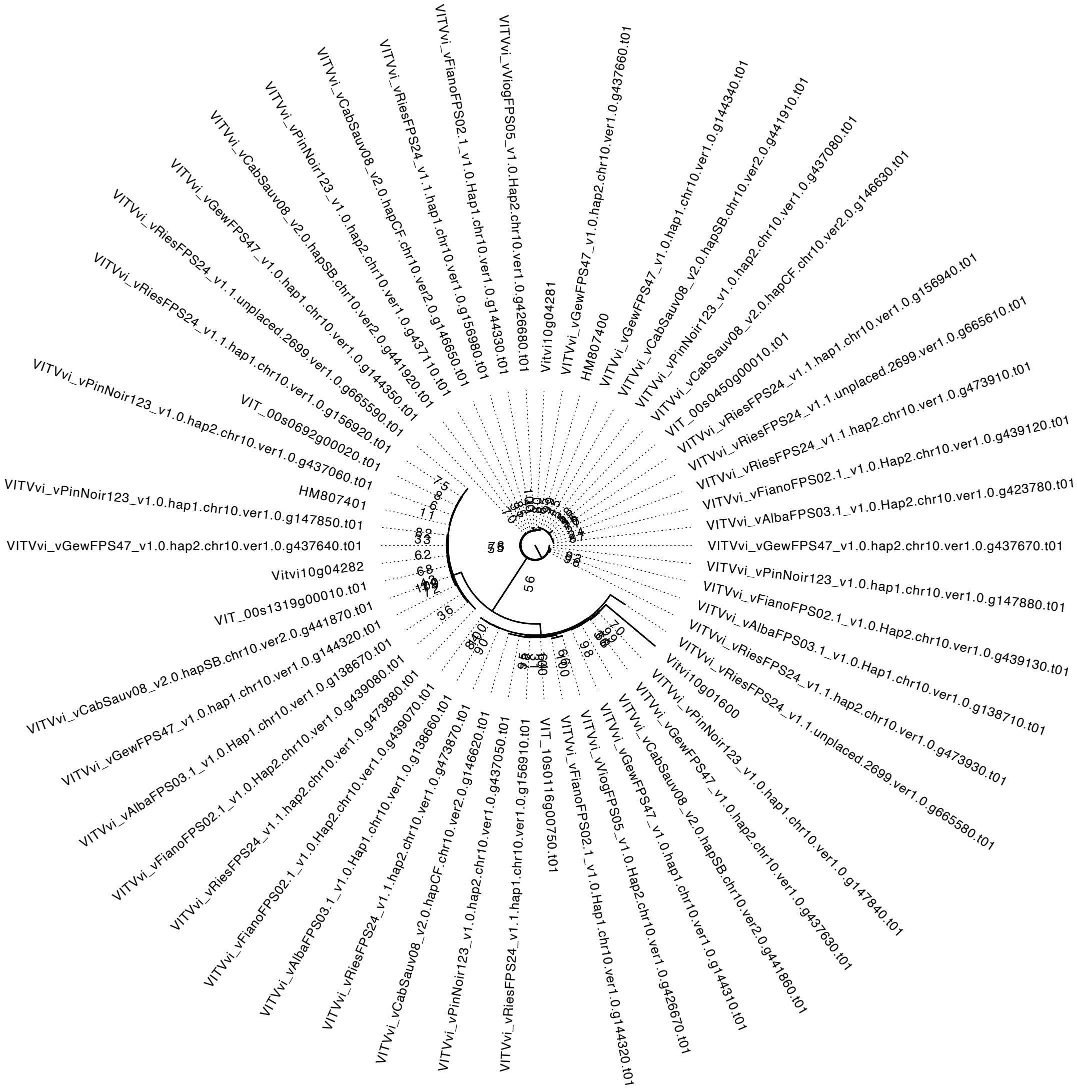

### Figure S6

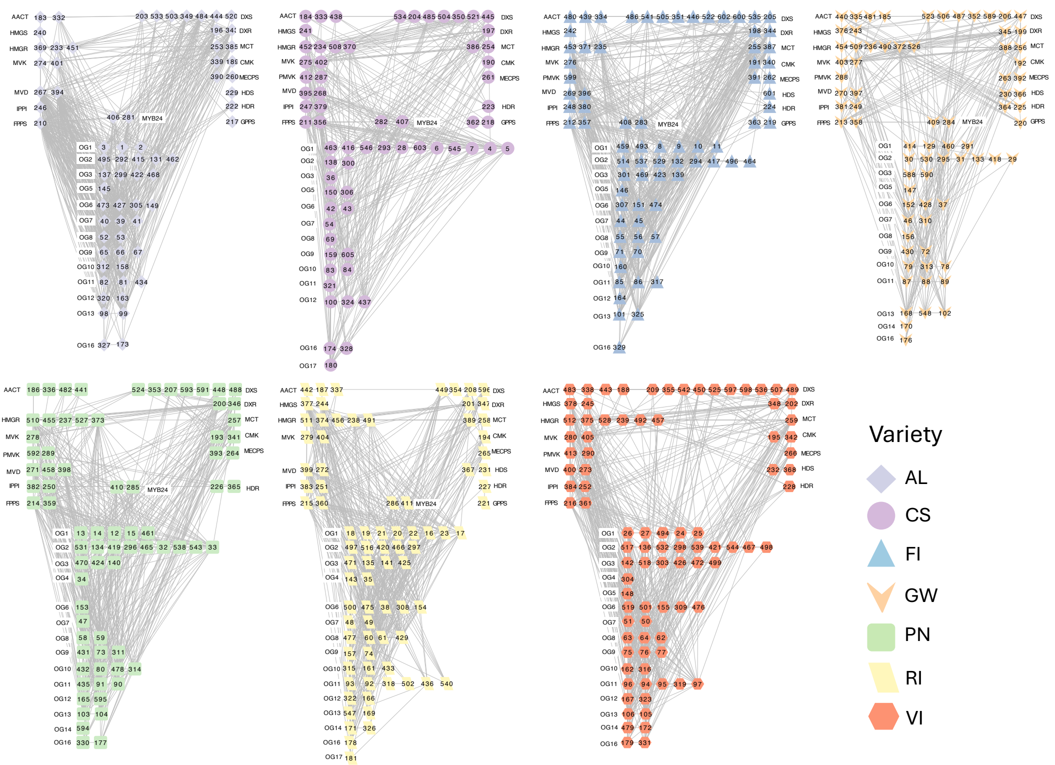

### Figure S7

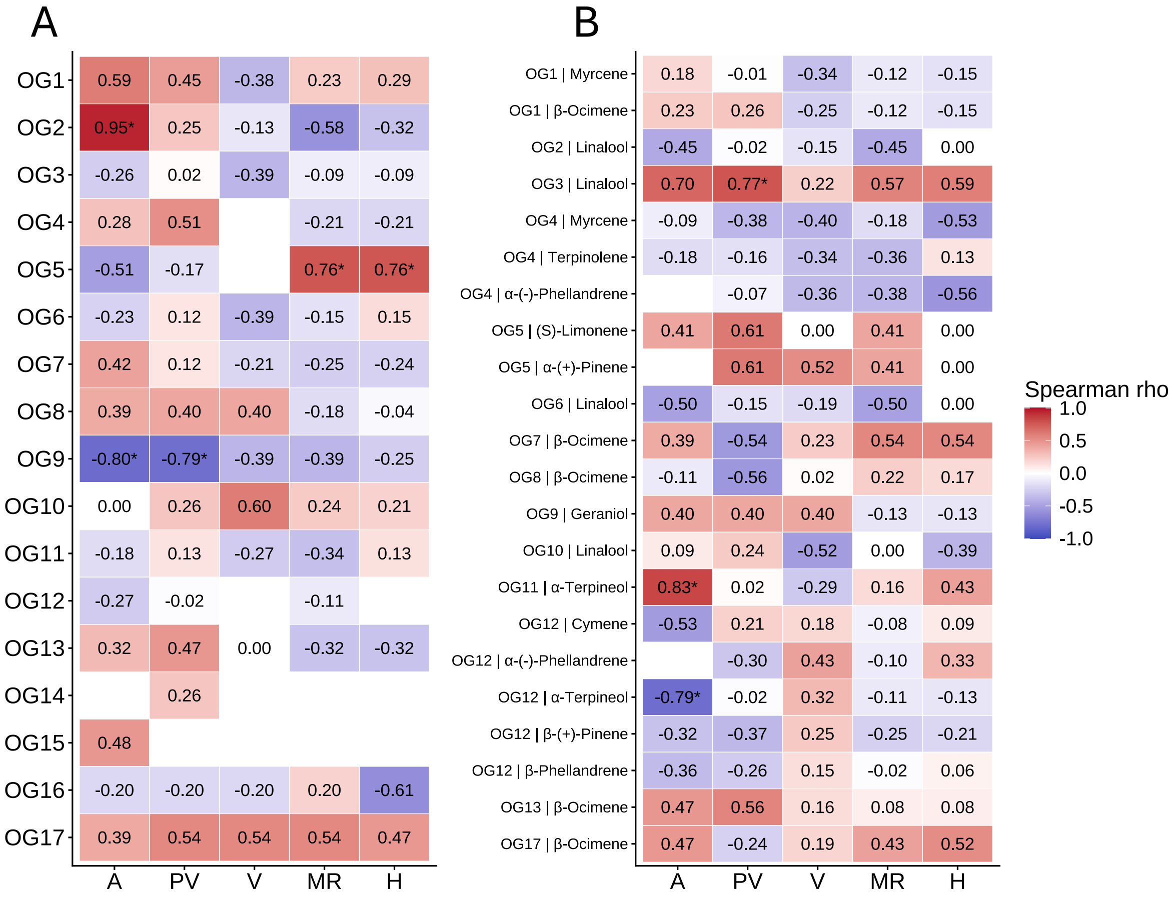
